# Volatility in coral cover erodes niche structure, but not diversity, in reef fish assemblages

**DOI:** 10.1101/2021.11.03.467170

**Authors:** Cheng-Han Tsai, Hugh PA Sweatman, Loïc M Thibaut, Sean R Connolly

**Affiliations:** College of Science and Engineering, James Cook University; Townsville, QLD 4811, Australia; Australian Institute of Marine Science; Townsville MC, QLD 4810, Australia; Australian Research Council Centre of Excellence for Coral Reef Studies; Townsville, QLD 4811, Australia; School of Mathematics and Statistics, University of New South Wales; Sydney, NSW 2052, Australia; Smithsonian Tropical Research Institute; Panama, Republic of Panama

## Abstract

Environmental fluctuations are becoming increasingly volatile in many ecosystems, highlighting the need to better understand how stochastic and deterministic processes shape patterns of commonness and rarity, particularly in high-diversity systems like coral reefs. Here, we analyze reef fish time-series across the Great Barrier Reef to show that approximately 75% of the variance in relative species abundance is attributable to deterministic, intrinsic species differences. Nevertheless, the relative importance of stochastic factors is markedly higher on reefs that have experienced stronger coral cover volatility. By contrast, α-diversity and species composition are independent of coral cover volatility but depend on environmental gradients. Our findings imply that increased environmental volatility on coral reefs erodes assemblage’s niche structure, an erosion that is not detectable from static measures of biodiversity.

**One-Sentence Summary:** Coral cover volatility modulates how stochastic and deterministic processes shape commonness and rarity in coral reef fishes.

## Main Text

Coral reef ecosystem dynamics and community structure are profoundly influenced by episodic, stochastic disturbances (*1–6*). Differences in species’ susceptibility to disturbances (*7, 8*), and differences in species’ rates of recovery from such disturbances (*9, 10*), suggest that environmental stochasticity can have a substantial role in shaping species’ commonness and rarity on coral reefs. However, the fossil record also provides ample evidence of persistent differences in species’ abundances over long time scales (*11, 12*), including consistently rare species (*11, 13*). This indicates that species’ intrinsic traits also influence their relative commonness or rarity. The combination of high diversity and pronounced stochasticity in species’ relative abundances has made coral reefs a model system for tests of neutral theory (*14–17*) and fluctuation-mediated theory of species coexistence (*2, 3*). Tests of neutral theory revealed that both reef coral and reef fish assemblages appear to have more heterogeneity in species’ relative abundance than that of neutral models can explain (*15, 17, 18*). Nevertheless, the fact that neutral theory is insufficient to explain community structure on coral reefs leaves its relative importance unresolved. Specifically, neutral ecological drift could explain most of the variation in species abundances, with species’ niche differences additionally driving small heterogeneity in abundances. Alternatively, neutral drift could be relatively unimportant, with most of the variation in species abundance driven by differences in the demographic characteristics of species. Moreover, the current widespread environmental disruption of reef community structure, function, and dynamics raises the possibility that the relative importance of these different classes of processes is changing, with potential consequences for the future of reef biodiversity.

Here, we use an approach derived from a stochastic-dynamic theory of community structure to partition the variance of the temporal dynamics of relative species abundances (*19*) and apply it to a unique, regional-scale, spatially replicated time series of species abundances of fishes on the Great Barrier Reef (see Materials and Methods and Fig. 1). Our analysis reveals that reef fish communities are structured disproportionately by persistent, intrinsic differences among species, rather than by stochastic fluctuations in population growth rates (Fig. 1E). Specifically, these differences explain a substantially larger proportion of variation in reef fish community structure (75% on average [95% CI: 72%-78%]), compared to stochastic fluctuations in population growth rates (18% on average [95% CI: 15%-20%]) (Fig. 1E). Only ∼7% of the variance is attributable to additional sources of variance, such as demographic and sampling variance (Fig. 1E). Despite the well-documented importance of episodic disturbances in coral reef ecology, our findings show that species differences underlie persistence in commonness versus rarity through time. This suggests that ecological traits of species influencing long-term mean abundances are disproportionately responsible for the variation in species’ abundances in reef fish assemblages.

**Fig. 1.**
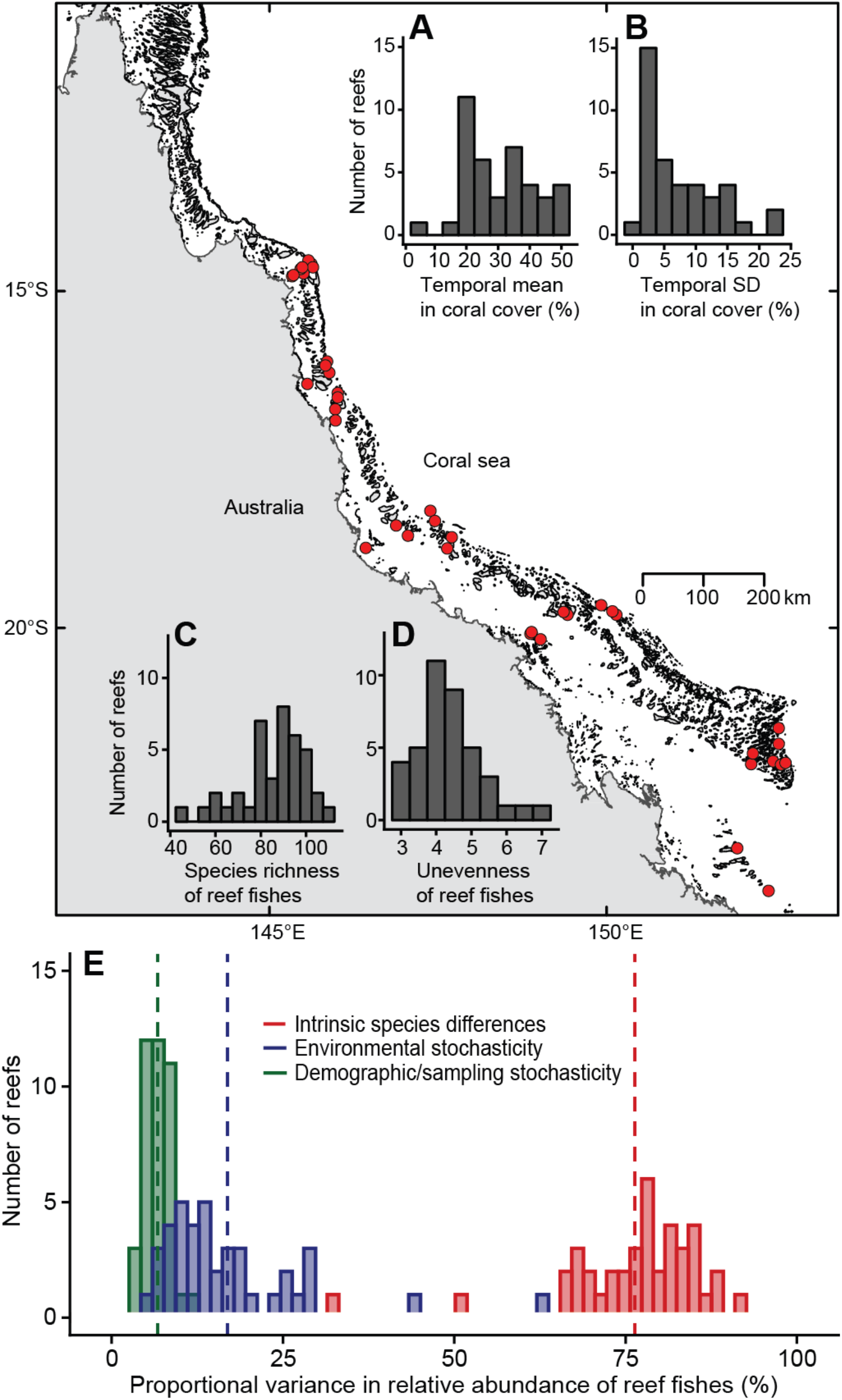
Map showing reefs included in all analyses, along with frequency distributions of explanatory and response variables. On the map, red circles show locations of the n=40 reefs used in this study. (**A**-**B**) Frequency distribution of the temporal mean and standard deviation in coral cover that are proxies for environmental volatility across study sites. (**C**-**D**) Frequency distribution of (time-averaged) species richness and unevenness of reef fish communities across study sites. (**E**) Frequency distribution of the proportional variance in relative species abundances attributable to deterministic intrinsic species differences that produce differences in long-term mean abundances (red bars; n=40), to environmentally induced stochastic fluctuations in species’ growth rates (blue bars; n=40), and to residual effects such as demographic and sampling variance (green bars; n=40). Color-coded, dashed lines indicate the mean value of the corresponding variance component.

Despite the lower relative importance of stochastic fluctuations in population growth rates as a driver of variation in commonness and rarity overall, there is substantial variation in the relative magnitudes of variance components among the reefs in our study (i.e., the spread in variance components; Fig. 1E), and much of this spatial variation is explainable (Fig. 2, Table S1). In particular, the relative importance of persistent species differences in driving reef fish abundances varies strongly with temporal volatility in coral cover (Fig. 2). Reef fish assemblages are less deterministically structured on reefs with more volatile coral cover, especially when mean coral cover is high (cf. Fig. 2A and 2C). The interactive effects of the mean and temporal standard deviation of coral cover together explain about 40% of the variation in these two variance components of fish species abundance (*R*^2^=0.39, *p*<0.001 for the variance component of persistent species differences, and *R*^2^=0.4, *p*<0.001 for the variance component of stochastic fluctuations; Table S1). Model selection favors this interactive model over all alternatives considered (Table S1). Surprisingly, latitude and cross-shelf position, which can serve as proxies for a range of regional-scale environmental gradients (Fig. S1), explained almost none of the spatial variation in the relative importance of persistent, intrinsic species differences (Table S1), despite the fact that reef fish species composition is known to change quite markedly from inshore to offshore (*20, 21*), and indeed cross-shelf position explains about 3–4 times more variation in community composition than coral cover variables in our data (Fig. S2).

**Fig. 2.**
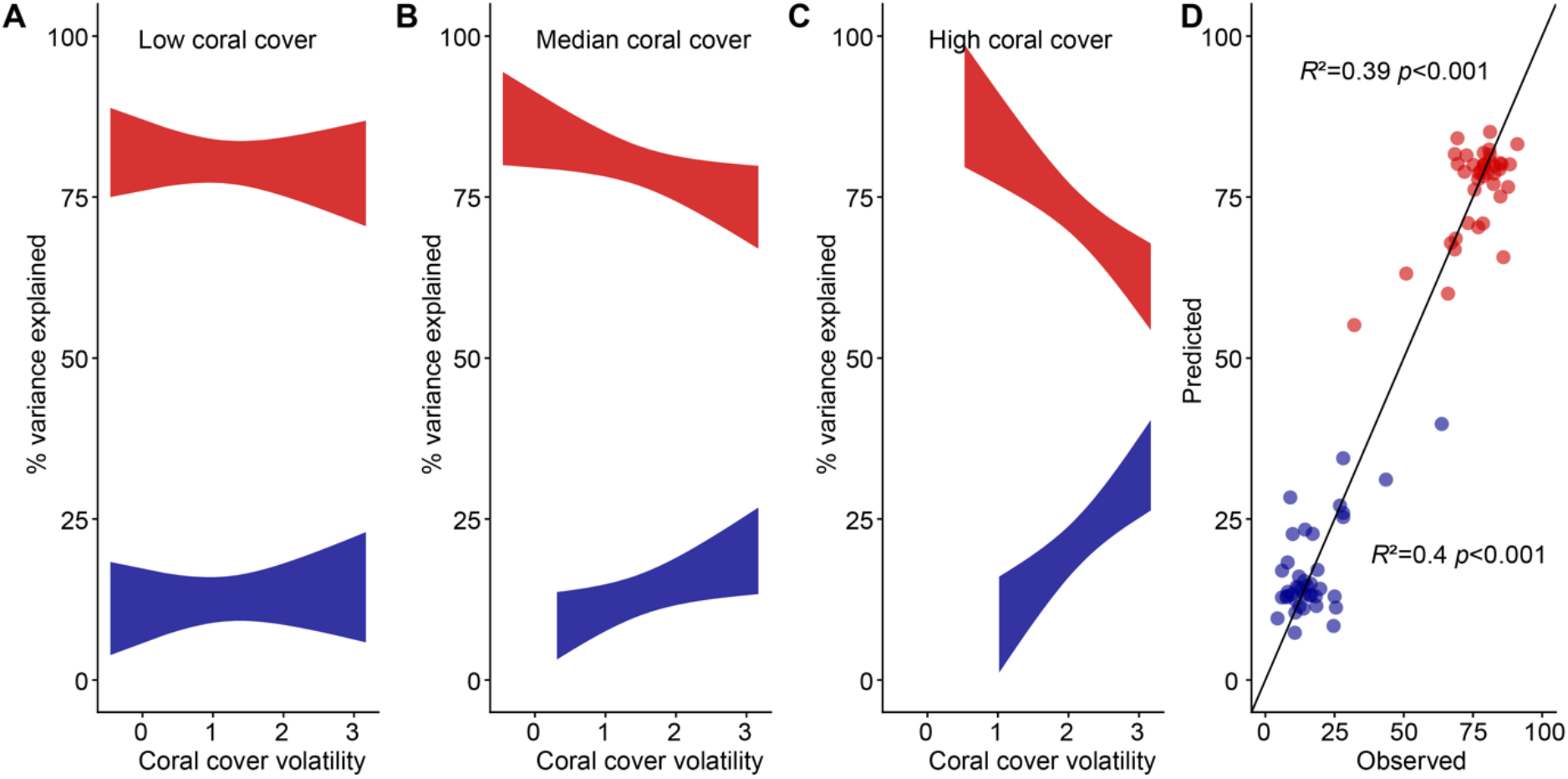
Deterministic and stochastic variance components in relative species abundance of reef fishes depend on coral cover volatility, but not on physical environmental gradients. (**A**-**C**) Relationships between the reef-scale coral cover variables (temporal standard deviation and mean of coral cover for each reef) and the relative importance of variance components in structuring fish species-abundances across reefs. The relationships are plotted using parameter estimates for the lowest-AIC models, with interactive effects of the temporal SD and mean of coral cover as explanatory variables, and variance components of fish community structure as response variables (n=40 reefs, Supplementary Table S1). The red bands represent the 95% C.I. of the proportional variance attributable to persistent, intrinsic species differences, while the blue band represents the 95% C.I. of the proportional variance attributable to environmental stochasticity. To illustrate the interactive relationships, the 1^st^, median and 3^rd^ quartiles of mean coral cover are fixed in panels (**A**), (**B**), and (**C**), respectively, and the fitted relationship between the natural logarithm of the standard deviation of coral cover and variance component values are plotted for the corresponding value of mean coral cover. (**D**) The relationship between observed and predicted values from the corresponding OLS regression models whose fits are plotted in panels (**A**-**C**). The solid line is the unity line (observed = predicted).

In contrast to the dynamical quantities represented by the variance components, coral cover volatility explains little variation in static measures of community structure (i.e., time-averaged richness and evenness; Methods) of the reef fish assemblages (*R*^2^∼0 in all cases; Table S1). Instead, these α-diversity quantities vary strongly and interactively with latitude and cross-shelf position, which together explain about 55% and 71% of the variation in richness and evenness, respectively (Fig. 3). Specifically, species richness increases and unevenness decreases (i.e., evenness increases) towards the equator, but the increases in richness and decreases in unevenness are much steeper on the inner shelf than the outer shelf of the Great Barrier Reef (Fig. 3). This might be due to the fact that human population density and associated coastal impacts disproportionately affect nearshore versus offshore reefs and also decrease towards the equator on the Great Barrier Reef (*22*). Alternatively, the interaction effect of cross-shelf position and latitude might be explained by natural covariation between locations and oceanographic conditions (*23*).

**Fig. 3.**
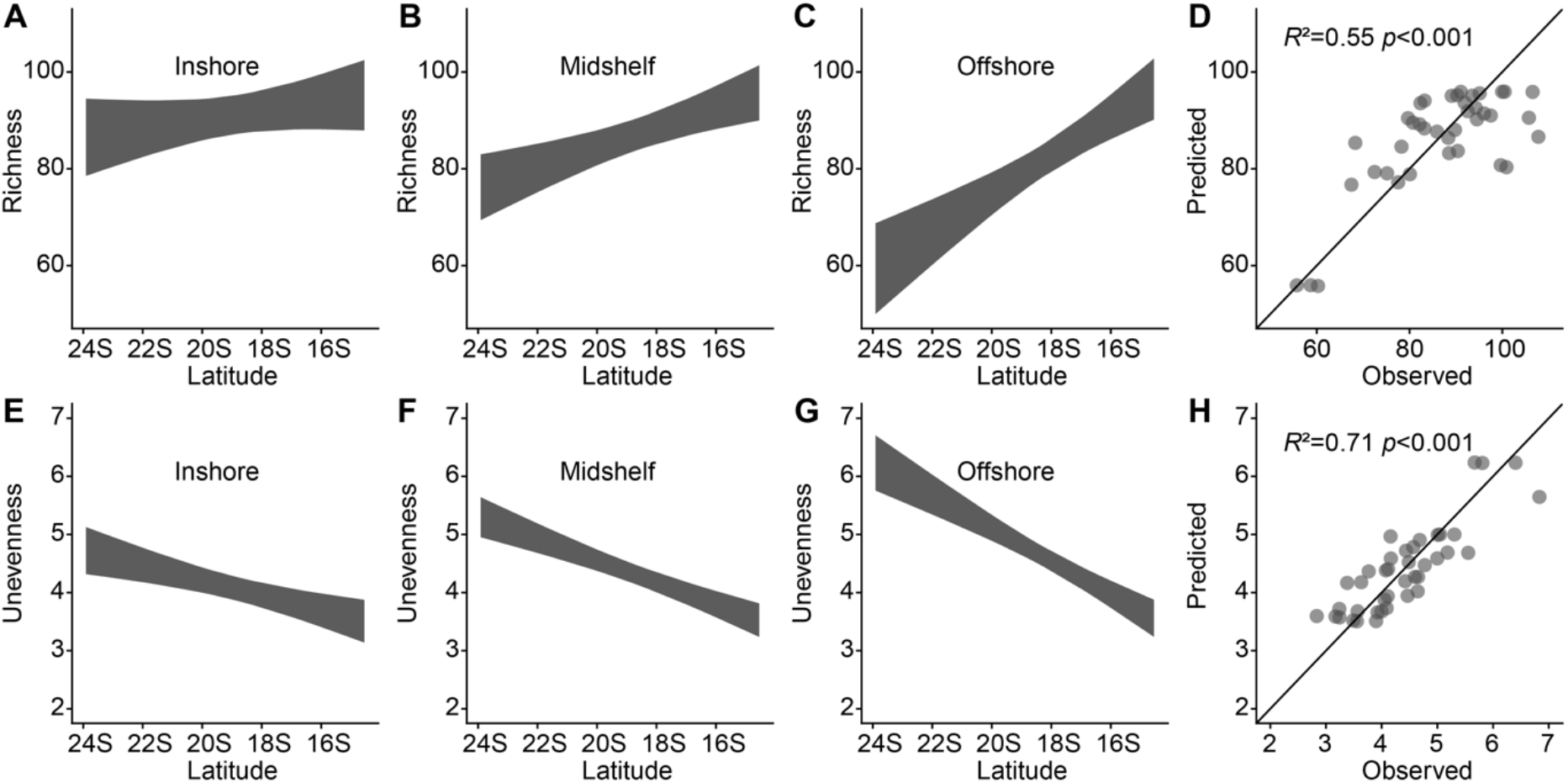
α-diversity (richness and unevenness) depend on physical environmental gradients, but not on coral cover volatility. (**A**-**C**) Relationship between time-averaged fish species richness and the interaction of latitude with cross-shelf position. (**E**-**G**) Relationship between time-averaged unevenness and the interaction of latitude with cross-shelf position. Grey bands are the 95% C.I. predicted from the lowest-AIC models for richness and unevenness (n=40 reefs, Supplementary Table S1). To better illustrate the interactive relationships, the 1^st^, median, and 3^rd^ quartiles of cross-shelf positions are fixed in panels (**A**, **E**), (**B**, **F**), and (**C**, **G**), respectively, and the relationship between richness or unevenness as a function of latitude are shown for the corresponding value of cross-shelf position. Note that values of the cross-shelf position increase from the offshore towards the coast. (**D**, **H**) The relationship between observed and predicted values from the OLS regression models corresponding to the fitted relationships in (**A**-**C**) and (**E**-**G**), respectively. The solid line is the unity line (observed = predicted).

We initially suspected that much of this heterogeneity in species’ abundances was a consequence of the functional diversity of the reef fish assemblage. However, extended analysis of our data indicated that this was not the case. After dividing all our fishes into functional groupings, we had sufficient species richness in three of our groups (herbivores, planktivores, and benthic invertivores) to repeat our entire analysis on each of them separately (see Materials and Methods, Table S2). We found that the proportion of variance in abundance attributable to persistent species differences within each of these functional groups was similar to the proportion for the fish fauna as a whole (78%, 83% and 75% on average for herbivores, planktivores, and benthic invertivores, respectively; Table S3-S5, Fig. S3). Moreover, the variance components of relative species abundance changed with coral cover volatility in a manner virtually identical to when the data were analyzed in the aggregate (Table S6, Fig. S4). Richness and evenness of functional groups also varied with respect to latitude and cross-shelf position similarly to the whole fauna, although in contrast to the variance components of community structure, the magnitude of richness and unevenness estimates varied substantially among the groups (Table S6, Fig. S5-S7).

The large proportion of variance in reef fish species’ abundances (including functional groups) attributable to persistent, intrinsic species differences underscores the importance of niche structure and places a clear upper bound on how much of the species abundances can be explained by neutral ecological drift (Fig. 1E and Fig. S3). This upper bound is provided by our estimates of deterministic niche structure (i.e., community determinism resulting from intrinsic species differences). Whereas species’ differential responses to environmental fluctuations (i.e., response diversity) likely also contribute to the stochastic-fluctuation variance component (Fig. 1E). Indeed, half of the variation among reefs in this variance component is explained by volatility in coral cover, suggesting that response diversity could also contribute to the variation in species’ abundances that are owing to stochastic fluctuations (Fig. 2). If so, the importance of niche structure may be higher than our deterministic estimates.

One of our key findings–that nearly half the variation in community determinism spanning 10 degrees of latitude across the Great Barrier Reef can be explained by just two explanatory variables linked to coral cover volatility–highlights the risk of ongoing erosion of niche structure among coral reef fishes (Fig. 1-2, Table S1). In this system, regional variation in coral cover dynamics during the time frame of our analysis has been driven substantially by episodic disturbances, including cyclones, crown-of-thorns starfish outbreaks, and coral bleaching (*4, 24, 25*). As climate change accelerates, coral bleaching is almost certain to overtake other disturbances as a key driver of increased coral cover volatility, if it has not already (*5, 26, 27*). Our findings suggest that such volatility will further erode the role of intrinsic fish species traits in structuring community abundance patterns, most dramatically on reefs with the highest levels of average coral cover (Fig. 2). Moreover, because the erosion in assemblage structure is not reflected in static measures of community structure, such as richness, evenness or composition (Fig. 3, Fig. S2), reliance on these metrics might fail to provide sufficient early warning for important changes in the processes structuring coral reefs ecosystems, and highlight the urgent need for long-term community-level abundance data to identify signs of ecological degradation (*6, 28, 29*).

## Acknowledgements

We thank members of ecological modelling group at James Cook University for discussions. We thank LTMP of Australian Institute of Marine Science for maintaining the data. We thank HPC of James Cook University for computing resources. C.-H. Tsai was supported by AIMS@JCU and ARC Centre of Excellence for Coral Reef Studies.

## Author contributions

C.-H.T., H.S. and S.C. devised the research program; H.S. supervised and provided the LTMP data; C.-H.T. performed analysis with help from L.T. and S.C.; C.-H.T. and S.C. wrote the first draft of manuscript, and all authors were involved in interpreting the results and contributed to the final draft of manuscript.

## Competing interests

The authors declare no competing financial interests.

## Data and materials availability

R code and data required for analyses are available in Supplementary Materials and at https://github.com/TsaiCH/simsEngenVPRSA. LTMP data is accessible by contacting H.S.

## Supplementary Materials

Materials and Methods

Figs. S1 to S8

Tables S1 to S10

Supplementary Text

Supplementary Simulation Study

References (*30–53*)

## Supplementary Materials

**This PDF file includes:**

Materials and Methods

Supplementary Text

Figs. S1 to S8

Tables S1 to S10

Supplementary Simulation Study

Simulation Study Figs. 1 to 6

References (*30–53*)

### Materials and Methods

#### Fish community data and environmental covariates on the Great Barrier Reef

We use data from the Australian Institute of Marine Science Long-Term Monitoring Program (LTMP), which has made visual surveys of benthos and fish communities on reefs spanning 10 degrees of latitude on the Great Barrier Reef for more than 20 years (*30*). The surveys themselves are hierarchically structured: 3 sites on the reef slope at approximately 6-9m depth were selected, usually on the NE faces of 40 reefs of our study (*30*). At each site, five permanently marked 5m × 50m transects were established for censusing all fishes other than small damselfishes, which were counted on 1m × 50m sections of the same transects. Transects were separated by about 10m. For these analyses, we used the community data from 1994 to 2004 (11 years) because this was the only interval during which each reef was surveyed annually (the frequency of surveys changed after the Great Barrier Reef Marine Park was rezoned in 2004).

The statistical analyses focus on counts of fish identified to species and percentage cover of live coral at each survey reef. Fish species are counted visually for a prescribed list of species representing 13 families: *Pomacentridae*, *Acanthuridae*, *Serranidae*, *Lutjanidae*, *Scaridae*, *Caesionidae*, *Chaetodontidae*, *Labridae*, *Lethrinidae*, *Haemulidae*, *Holocentridae*, *Siganidae*, and *Zanclidae*. All species examined here are largely non-cryptic and easily identified underwater, and thus cryptic species groups, such as gobies, were excluded. A full list of species observed each year are included in the appendices of each LTMP status report (*30*) (also see Table S2 for species list used for our analysis). Corals were identified to relatively broad taxonomic and morphological categories, but we consider only total hard coral cover in our analyses. We pool fish community and coral cover data at the scale of the entire reef, summing abundances over all 15 transects surveyed at each reef. Percentage cover is similarly averaged across transects and sites within reefs. We adopt this approach to reduce stochastic sampling error, thereby obtaining more precise estimates of the community structure statistics that are of interest in this study.

Because the small-sized fish taxa (mainly *Pomacentridae*) were surveyed in narrower transects than other larger fish taxa, we used subsampling to rescale the abundances of large-sized species to standardize sampling effort. Each fish counted on the wider transects was given a 20% probability of appearing in the sub-sample (because the small-fish transects covered only 20% of the area of large-fish transects). It is these sub-sampled data that were used for our analyses.

For each reef, we extracted the temporal average (11-yr mean), standard deviation (SD) and coefficient of variation (CV) in coral cover as proxies for disturbance-induced coral cover volatility. We also extracted each reef’s latitude and cross-shelf position, where latitude was measured by degrees from the equator and cross-shelf position was the standardized distance to the nearest continental shelf boundary (i.e., 0 represents the shelf boundary and 1 represents the coast, respectively). We used latitude and cross-shelf position as proxies for major environmental gradients because community structure on the Great Barrier Reef is known to vary strongly along both gradients, and because they are strongly correlated with environmental variables, such as mean and variability in temperature, and variation in terrestrial runoff, which are known to influence community structure (Fig. S1).

#### Partitioning variance in relative species abundance: theoretical framework

We used the partitioning approach of Engen et al. (2002) (*31–33*) to quantify the contribution of deterministic species differences, relative to environmental and demographic stochasticity in driving the total variance in relative species abundances (hereafter, variance partitioning of relative species abundance, VRPSA). These variance components can be estimated from how the correlation in a community’s log-abundances decays over time, i.e., the temporal autocorrelation in relative species abundance:

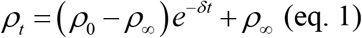

where *ρ_t_* represents the correlation coefficient of log species-abundances of a community at time lag *t* (i.e., it is a measure of community similarity between species’ log-abundances in two different years). This quantity is modelled as an exponential function of time lag *t*. That is, for all pairs of years on a reef, we estimated the correlation coefficient between these two samples, and then analyzed how the strength of this correlation decreased as a function of the time elapsed between the two samples (here, time lag or interval between two samples ranges from 1 to 10 years). Parameter *ρ*_∞_ represents asymptotic similarity. For a community in the absence of persistent niche structure in abundance (e.g., where all species have the same mean abundance, and variation in species abundances is due entirely to their different responses to environmental fluctuations and independent demographic stochasticity), *ρ*_∞_ would be zero. For a community in which environmental fluctuations play no role, *ρ*_∞_ would be large (and in the absence of demographic or sampling stochasticity, it would tend to unity). Parameter *ρ*_0_ is the intercept (i.e., the expected correlation in species’ log-abundances for a time lag of zero), and thus 1-ρ_0_ represents variation that not captured by the other two components (including demographic variance, as well as sampling effects due to local aggregation). *δ* measures the strength of density-regulation in the system: it would be larger in assemblages that revert quickly toward their long-term mean relative abundances after a disturbance (see Supplementary Text for further explanation of eq. 1).

The correlation coefficients *ρ_t_* were estimated by fitting the bivariate Poisson-lognormal distribution to all possible pairs of surveys at each site, and these correlation coefficients were then modelled as a function of the amount of time separating the samples following eq. 1. This distribution assumes that the two surveys represent random samples of individuals from two communities whose species abundances follow Poisson-lognormal distributions with correlation coefficient *ρ_t_* That is, the correlation coefficient parameter estimates the true underlying correlation in species’ log-abundances between the communities at the two sampling times, taking account of the fact that each abundance distribution in the data represents an incomplete sample from the community, and, therefore, this quantity can be conceptualized as a measure of temporal beta-diversity (*32, 33*). This model is justified because the static species-abundance distributions of these reef fish assemblages have been shown to be well-described by the Poisson-lognormal distribution (*34*). As a further check, we conducted parametric bootstrap tests (Bootstrap N=100 for each fitted bivariate Poisson-lognormal distribution) to verify that the bivariate Poisson-lognormal is an adequate distribution for the LTMP data, and we found that none of our study reefs were statistically distinguishable from this distribution.

After fitting eq. 1 to the estimates of pairwise correlation coefficients as a function of time lag, variance components of relative species abundance can be obtained as follows:

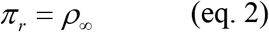

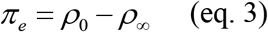

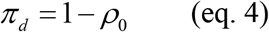

where *π_r_* represents the proportional variation in relative abundance that is due to persistent, deterministic differences among species, *π_e_* represents the proportional variation in relative abundance due to species’ responses to stochastic fluctuations in population size, and *π_d_* represents the proportional (residual) variation in relative abundance due to other processes, such as overdispersion.

#### Stochastic community dynamics model of VPRSA

Originally, the partitioning approach described above was explicitly derived from a stochastic theory of community dynamics. This theory characterizes the temporal dynamics of abundance of *S* species (richness is *S*) according to the stochastic Ornstein-Ühlenbeck process:

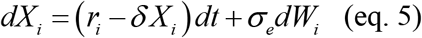

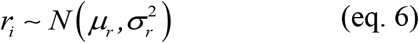

where *X_i_* represents the abundance of species *i* on a logarithmic scale, *r_i_* is the intrinsic population growth rate of species *i*, *σ* measures the strength of density dependence, *σ_e_* scales the magnitude of environmental fluctuations in the growth rate (i.e., larger *σ _e_* implies larger fluctuations), and *dW_i_* models the fluctuations themselves as a Brownian (Wiener) process. Eq. 6 specifies that intrinsic growth rates, *r_i_*, vary among species according to a normal distribution with mean *μ_r_* and variance *σ* ^2^. Eq. 5 can be interpreted as a continuous-time analogue of discrete-time Gompertz-type community dynamics.

Analysis of the model in eqns 5-6 shows that each species’ abundance fluctuates around a species-specific equilibrium “carrying capacity”, 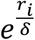, and both the carrying capacities, and the abundances themselves, follow lognormal distributions among species. Notably, the stationary distribution of species’ abundances in the community remains lognormal, even in the presence of some violations of the model’s simplifying assumptions, such as the incorporation of correlations in species’ responses to environmental fluctuations, and of inter-specific interactions (*31, 32*) The model is therefore consistent with the reef fish data in this study whose species-abundance distributions are well-described as discrete, random samples of individuals from lognormal abundance distributions (i.e., Poisson-lognormal distributions). In addition, previous work suggests that the Gompertz form of density dependence is appropriate for these data (*35*). For models of eqs 5-6, the total variance in relative species log-abundance (hereafter 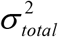) can be analytically partitioned into additive components as follows:

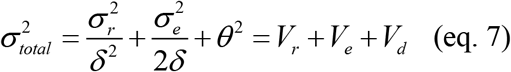

and thus the proportional variance components *π_r_*, *π_e_*, and *π_d_* would be equal to 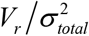, 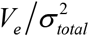 and 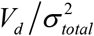, respectively. Extension of the model in eq (5) to explicitly incorporate demographic variance (intra-population heterogeneity in demographic rates and demographic stochasticity) suggested that such effects will be principally captured by the *πd* (and thus 1-*p*0) term, given that, like sampling effects, demographic variance will disproportionately affect observed abundances of the rare species (*31, 36*). However, to be conservative, our interpretation of our findings takes account of the possibility that part of the “environmental fluctuations” component of species-abundance variance could be due to such processes (including neutral ecological drift) (*37*).

Some of the assumptions of the stochastic dynamics model in eq. 5 are restrictive. Notably, it assumes that the strength of density regulation and the magnitude of environmentally induced fluctuations (i.e., proportional variance in population growth rate due to environmental fluctuations) are the same for all species, and it also assumes that there are no species interactions, nor covariation in species’ responses to environmental fluctuations. All of these assumptions might be violated to some degree in our study system. For a community showing Gompertz-type dynamics, the lognormal stationary distribution of abundances is quite robust to these simplifying assumptions. However, the robustness of variance component estimates has not yet been investigated previously, so we investigate this robustness in considerable detail in the Supplementary Simulation Study.

#### Estimating richness and unevenness based on VPRSA

We also use the Poisson-lognormal distribution to estimate (time-averaged) α-diversity measures of richness and unevenness of reef fishes as follows. We fit this distribution to each of the 440 species-abundance distributions (40 reefs × 11 years), using the method of maximum likelihood. Fitting this distribution yields maximum likelihood estimates of the standard deviation of log-abundance *σ̂* for the underlying community from which the data are a sample The skewness of a lognormal distribution is a function of only this parameter, implying that the skewness (i.e., the unevenness) of abundances in the fish community from which the data are a sample is a monotonically increasing function of *σ̂* (i.e., larger values indicate more uneven communities) (*32, 33*). This can be understood as a generalization of the evenness metric based on the variance in log abundances among species that accounts for the effects of incomplete sampling (i.e., the variance parameter is independent of sample completeness) (*33, 38*). Additionally, Poisson-lognormal fits can be used to estimate the total number of species in the community, by producing an estimate of the probability that a species is present in the community but not observed in the sample, *p̂*_0_. An estimate of total community richness is simply the number of observed species divided by 1− *p̂*_0_. We estimated the σ^2^ parameter of the Poisson lognormal distribution (i.e., the variance of the species abundance distribution from which the sample was drawn) for each year, and then we calculated the mean of these values over years for each reef as our reef-scale measure of unevenness (i.e., averaged σ^2^ is eqivalent to 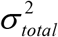 in eq. 7). Similarly, we used the mean of the estimated total community richness across years for each reef as our reef-scale measure of species richness.

#### Parameter estimation for VPRSA

To estimate the three variance components that completely partition the variation in relative abundances of fishes over years for each reef, we fitted a mixed-effects version of eq. 1 to the LTMP data set (Supplementary Text). Specifically, we fitted a family of nonlinear mixed-effects models, in which each of the fitted parameters (*^ρ^*^∞^, *^ρ^*^0^, *^δ^*) may be fixed constants for all reefs or they may randomly vary among reefs according to latent lognormal distributions (Supplementary Text). These models were parameterized in Template Model Builder (TMB) in R. We fitted models with different combinations of the three temporal autocorrelation function parameters (eq. 1) as fixed versus random, and we ranked model fits by AIC and bootstrapped AIC (Supplementary Text). We then checked for numerical stability of the model fits (i.e., we confirmed that the model’s random effects parameter estimates were valid), and we chose the best-fitting model (by AIC and bootstrapped AIC) that yielded a numerically stable fit as our basis for inference (Supplementary Text). Then, we used the estimates of fixed and reef-level random effects of the best-fitting model to calculate variance components for each reef according to eqs 2-4.

#### Predicting among-reef variation in community properties obtained by VPRSA

We used ordinary least-squares (OLS) regression to investigate the extent to which variation among reefs in coral cover fluctuations (mean, standard deviation [SD] and coefficient of variation [CV] of coral cover at each reef), latitude, and cross-shelf position explain variation in fish community structure (variance components of community structure, species richness, and evenness). We chose latitude and cross-shelf position as our abiotic explanatory variables because many factors likely to affect community structure on coral reefs, such as temperature, turbidity, and nutrient loading vary strongly with one or both of these spatial variables (Fig. S1). SD and CV of coral cover were both right-skewed, so they were log-transformed to reduce the heterogeneity of leverage values associated with the different reefs in our analysis (model selection yielded the same best-fitting models when they were untransformed, and *R*^2^ values were higher, making our results conservative with respect to the conclusions drawn). Log(SD) and log(CV) of coral cover were very strongly collinear (*r* ∼ 0.9), and model selection always preferred models using SD rather than CV when analogous models were compared (e.g., AIC favors a multiple regression model with interactive effects of log(SD) and mean coral cover over a model with interactive effects of log(CV) and mean coral cover), so we have not presented results for the models using log(CV). None of the remaining explanatory variables were highly correlated with one another (Supplementary Table S7-S8). We used AIC and adjusted *R*^2^ for model comparisons. We chose the parsimonious model with fewer parameters (effects) whenever AIC differences among candidates were smaller than 2 because, in such cases, adjusted *R*^2^ favored the simpler models, and because the additional variables included in the more complex models tended to have comparatively weak effects. In practice, this affected only model selection for unevenness, where the lowest AIC model included an additive effect of mean coral cover (in addition to a latitude × cross-shelf position interaction) that was small in magnitude and led to a marginally worse adjusted *R*^2^ (Table S9).

Lastly, because the reef-level proportional variance components were estimated from fits of another statistical model, rather than being directly observed, we also performed a sensitivity analysis to test the robustness of our results to parameter uncertainty. Specifically, we estimated the uncertainty in our estimates of proportional variances by parametric bootstrap, resampling random effects values for the LTMP’s 40 reefs 10000 times from the variance-covariance matrix of the fitted nonlinear mixed-effects model based on TMB. Then, each bootstrap set of reef-level proportional variance component values were re-analyzed using the OLS regressions repeated on each of the bootstrapped data sets. We evaluated the robustness of our model selection procedure by quantifying the percentage of bootstrapped data sets for which each model was selected as the best model by AIC.

#### Sensitivity analysis of community properties among trophic/functional groups

Reef fish community data were separated into five major trophic/functional groups: herbivores, planktivores, piscivores, corallivores and benthic invertivores. These trophic groups were classified according to previously published functional classifications of reef fishes, supplemented by communication with experts in the field (Table S2). Of these functional groups, only three (i.e., herbivores, planktivores, and benthic invertivores) were sufficiently species-rich to analyze separately. We repeated all of the analyses conducted on the overall dataset, as described above, for each of the groups. We then compared the distribution of reef-scale estimates of variance components for the three trophic groups with each other, and with those for the mixed (i.e., the whole data, regardless of functional groups) assemblages, to determine whether the magnitudes of the variance components changed markedly when functional groups were considered independently. Formally, these comparisons were made with paired *t*-tests with Bonferroni correction for multiple comparisons.

To examine the potential influence of coral cover volatility on variance components of relative species abundance within functional groups, we fitted relationships between variance components and coral cover volatility using OLS regression. In the full model, temporal mean and standard deviation in coral cover, and trophic group were considered as explanatory variables. AIC was used for model selection, beginning with a model including all main effects and interactions. As above mentioned, effects of latitude, cross-shelf position and trophic group on richness and evenness were also examined in a similar fashion.

### Supplementary Text

To choose the optimal mixed-effects structure (with reefs as random effects) for fitting the autocorrelation function (i.e., eq. 1 in the main text), we first confirmed numerical stability of the random-effect estimates (to verify that we had a valid model), by fitting the models using both importance sampling and the Laplace approximation, both implemented in Template Model Builder (TMB) in R. Then, for those models that produced stable, valid parameter estimates, we used model selection by AIC to choose the best model (Table S10). To confirm our model selection results, we also calculated marginal AIC, a statistic that uses penalized marginal likelihood by bootstrap resampling (5000 bootstrap replicates in our case); this statistic has been proposed as a more robust model selection statistic than AIC for models that have a common fixed effect structure but differ in their random effects (*39, 40*).

The model with random effects on all three parameters (ρ_0_, ρ_∞_, δ) was highly numerically unstable, as shown by the poor agreement between random effects estimates produced by importance sampling and the Laplace approximation (*R*^2^<0.5 to the unity line for all three parameters) (Fig. S8 A-C). For comparison, our estimated best model, which contains random effects on ρ_∞_ and ρ_0_ but not δ, was numerically stable, shown by virtually identical random effects estimates produced by the two fitting approaches (*R*^2^>0.95 in every case) (Fig. S8 D-E). Of all possible combinations of random effects structures, only models including a random effect of ρ_∞_ were numerically stable, and all numerically stable models produced very similar parameter estimates (Table S10). We therefore used the fixed and random effect estimates from our best-fitting model of all models producing numerically stable fits in all subsequent analyses throughout this study.

### Supplementary Figs. S1 to S8

**Fig. S1.**
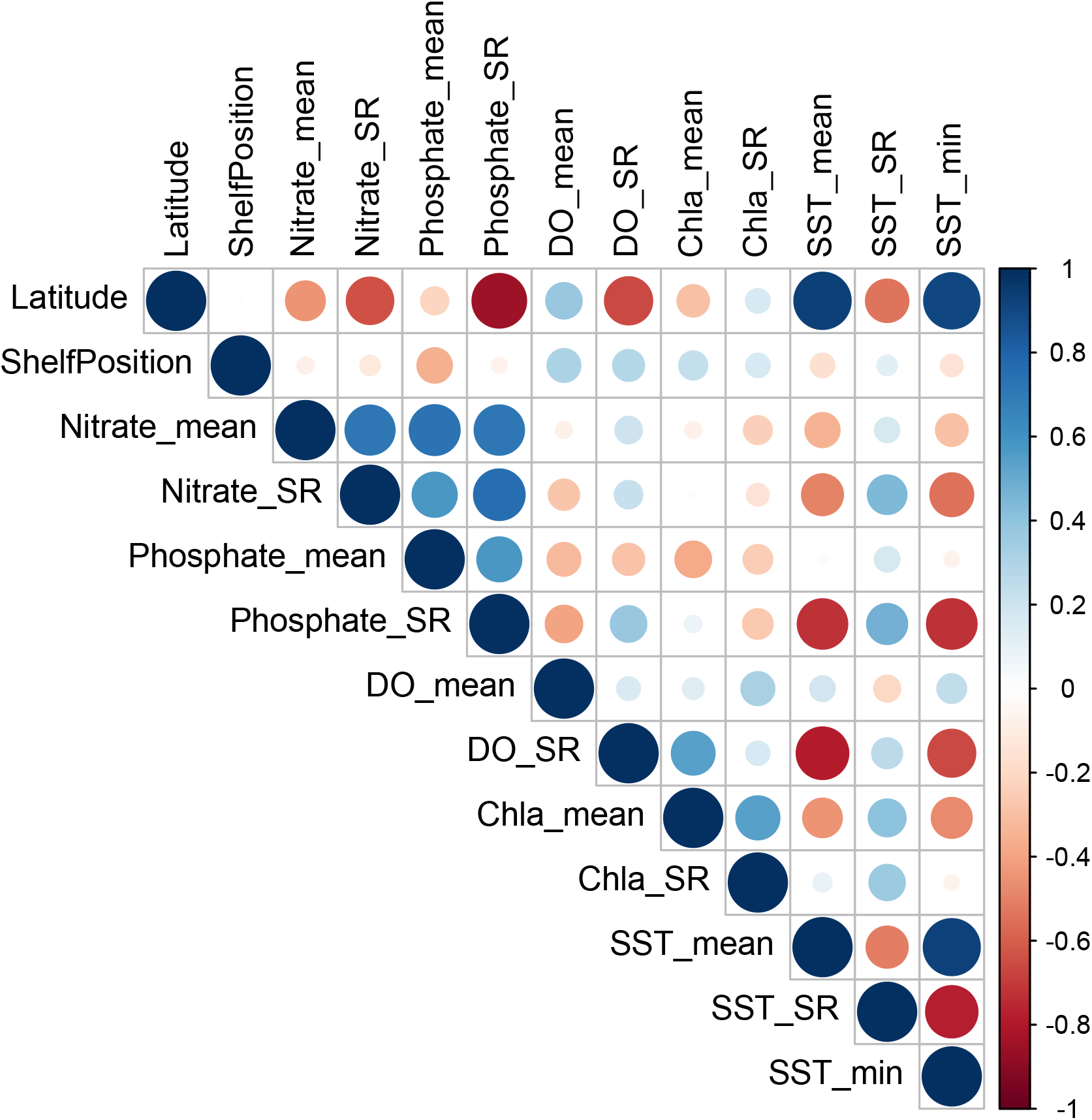
Latitude and cross-shelf position covary with oceanographic environmental conditions on the Great Barrier Reef. Pairwise Pearson correlations are presented in colour scale (n=40 reefs). ‘Nitrate_mean’, ‘Phosphate_mean’, ‘DO_mean’, ‘Chla_mean’, and ‘SST_mean’ represent long-term average of nitrate, phosphate, dissolved oxygen, chlorophyll-a, and sea surface temperature, respectively. Similarly, variables with ‘_SR’ and ‘_min’ attached represent averaged seasonal range and long-term minimum of the variables, respectively. Oceanographic environmental data on the Great Barrier Reef were taken from Matthews et al. 2019 (*41*).

**Fig. S2.**
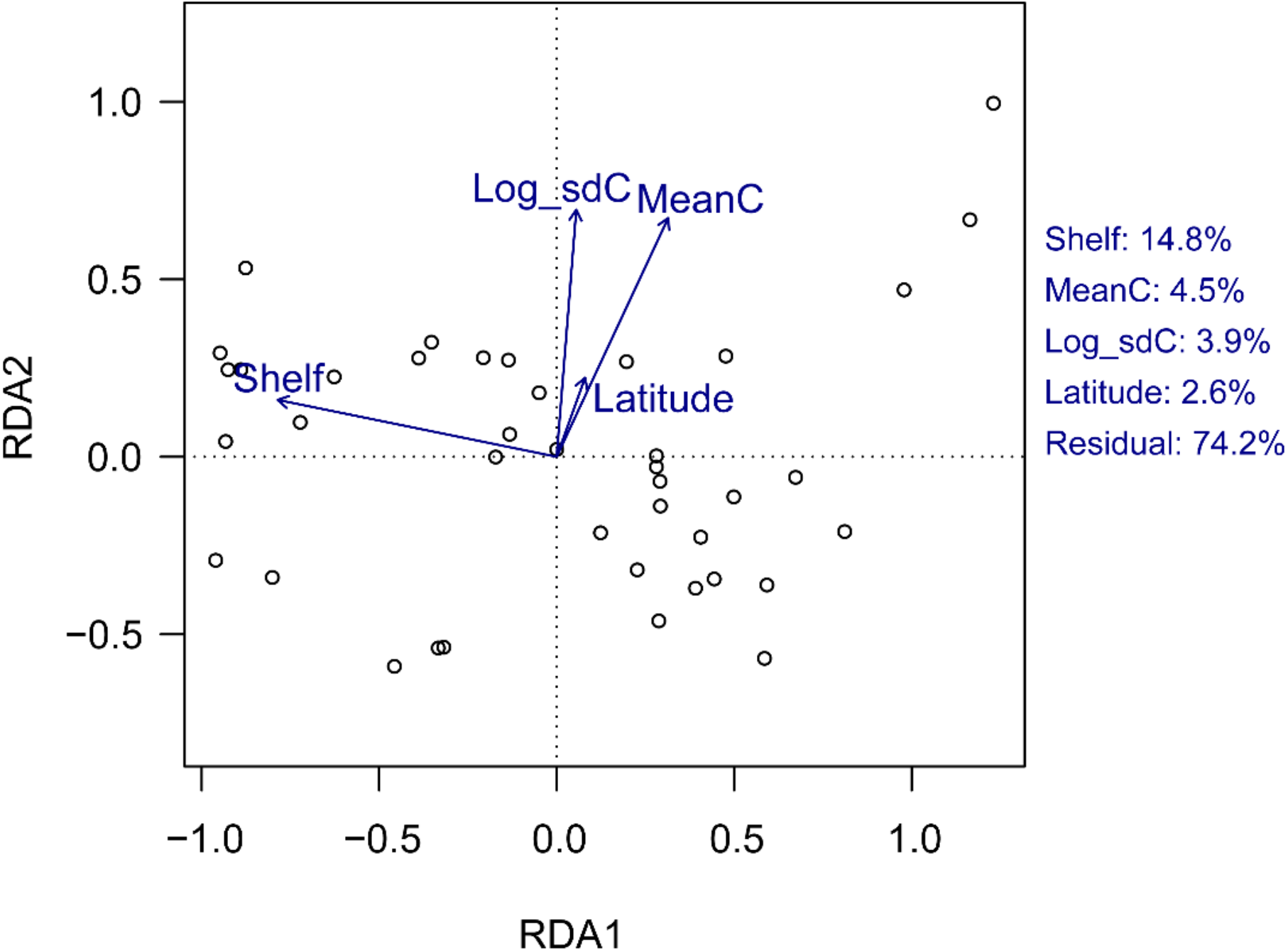
Reef fish community composition explained by the explanatory variables employed in our analyses. Redundancy analysis (RDA) constrained by environmental variables was used for the multivariate ordination plot. Open circles represent fish assemblages on coral reefs (n=40 reefs), and vectors represent major environmental variables, where ‘Shelf’, ‘MeanC’, ‘Log_sdC’ and ‘Latitude’ represents shelf position, average coral cover, coral cover volatility and latitude, respectively. On the right-hand side, the inertia (i.e., overall variance) is decomposed into RDA variance components explained by environmental variables in percentage. Reef fish community matrices are standardized prior to analysis. The RDA presented here uses abundance-weighted species composition, but the pattern is consistent with another RDA computed using only presence-absence data (not shown).

**Fig. S3.**
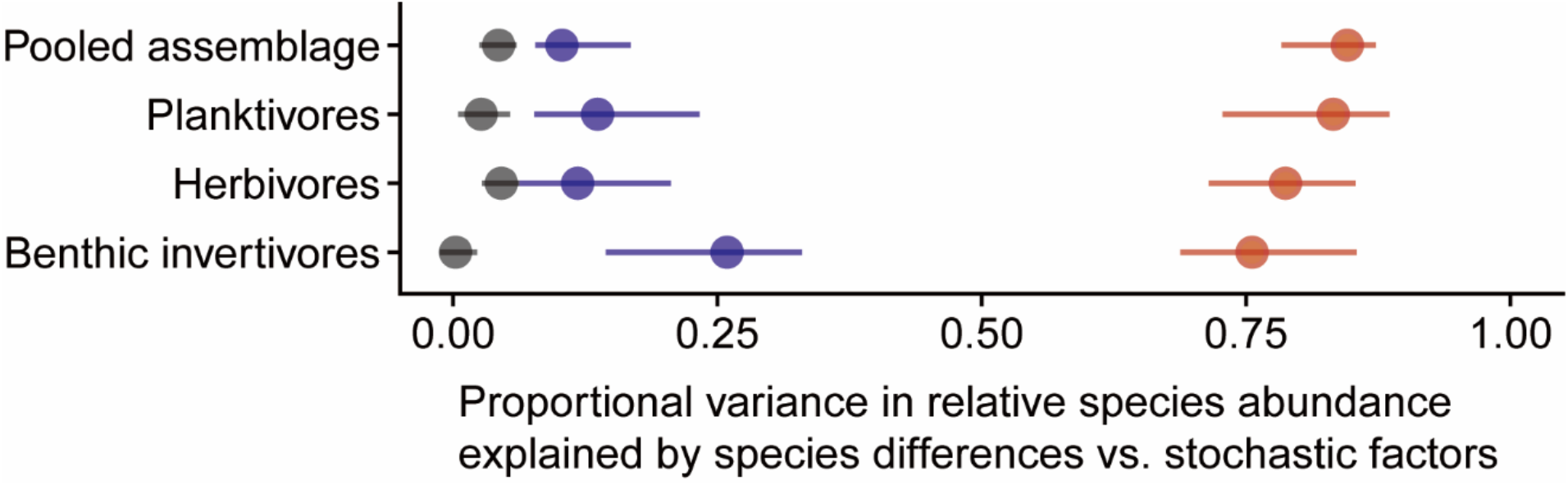
Estimated variance components of reef fish functional groups compared to pooled assemblage. Points and horizontal lines represent the median and the 1^st^ and 3^rd^ quantiles of estimates (n=40 reefs). Red color represents estimated variance component due to persistent species differences. Blue color represents estimated variance component driven by stochastic fluctuations in population growth rates. Grey color represents estimated variance component driven by demographic and sampling variance (or overdispersion). Variance components for the pooled assemblage are the same as in Fig. 1E in the main text and are presented here simply to facilitate comparison.

**Fig. S4.**
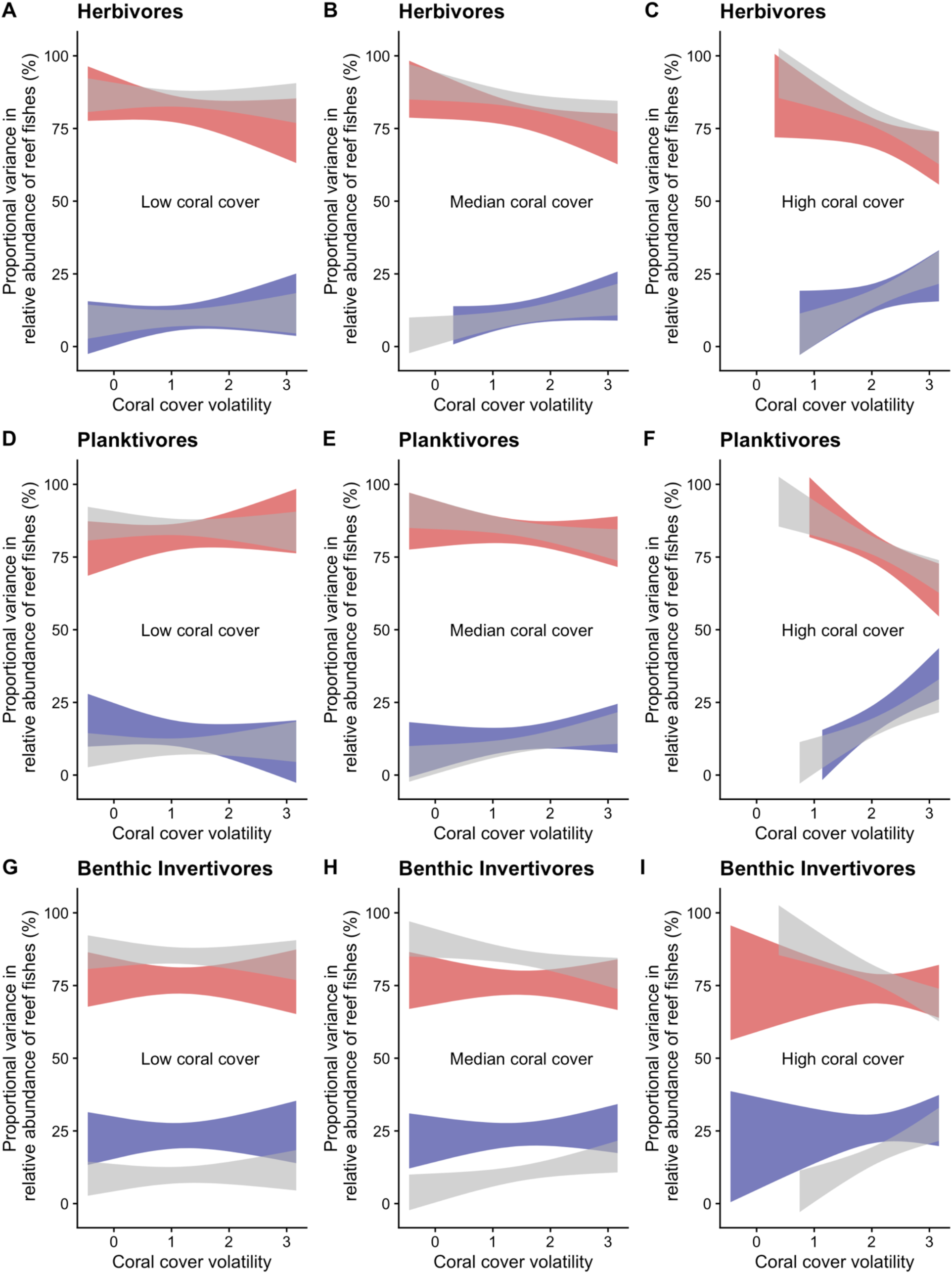
Relationship between coral cover dynamics and variance components of community structure of reef fish functional groups. Relationships between the reef-scale coral cover variables (temporal SD and mean of coral cover for each reef) and the relative importance of variance components in structuring species-abundances of fish trophic groups across reefs (n=40 reefs). (**A**-**C**) Herbivores. (**D**-**F**) Planktivores. (**G**-**I**) Benthic invertivores. The relationships are plotted using the lowest-AIC models, with interactive effects of the mean and SD of coral cover as explanatory variables, on variance components of fish community structure as response variables (Table S6). The red band represents the 95% C.I. of the proportional variance attributable to persistent species/niche differences, while the blue band represents the 95% C.I. of the proportional variance attributable to environmental stochasticity. For comparison, the grey bands represent the variance components for the original model, with functional groups pooled (as Fig. 2 in the main text). To illustrate the interactive relationships, the 1^st^, median and 3^rd^ quartiles of mean coral cover are fixed in panels (**A**-**C**), (**D**-**F**), and (**G**-**I**), respectively, and the relationship between the natural logarithm of the standard deviation of coral cover and variance component values plotted for the corresponding value of mean coral cover.

**Fig. S5.**
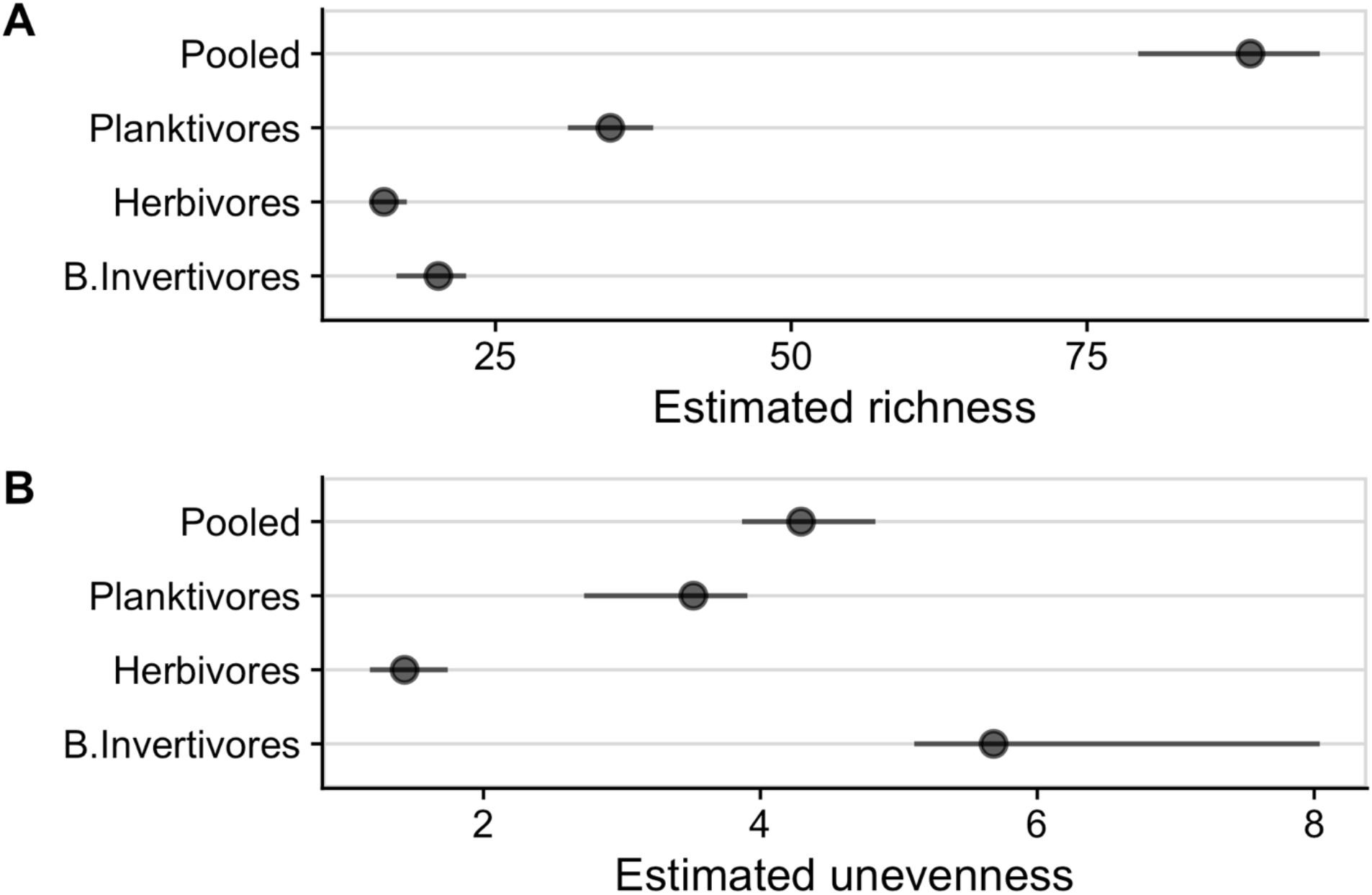
Estimated richness and unevenness of reef fish functional groups on the Great Barrier Reef. Points and horizontal lines represent the median and the 1^st^ and 3^rd^ quantiles of estimates (n=40 reefs). (**A**) Estimated species richness, and (**B**) estimated unevenness of trophic groups, compared to those of pooled assemblage presented in Fig. 1 in the main text.

**Fig. S6.**
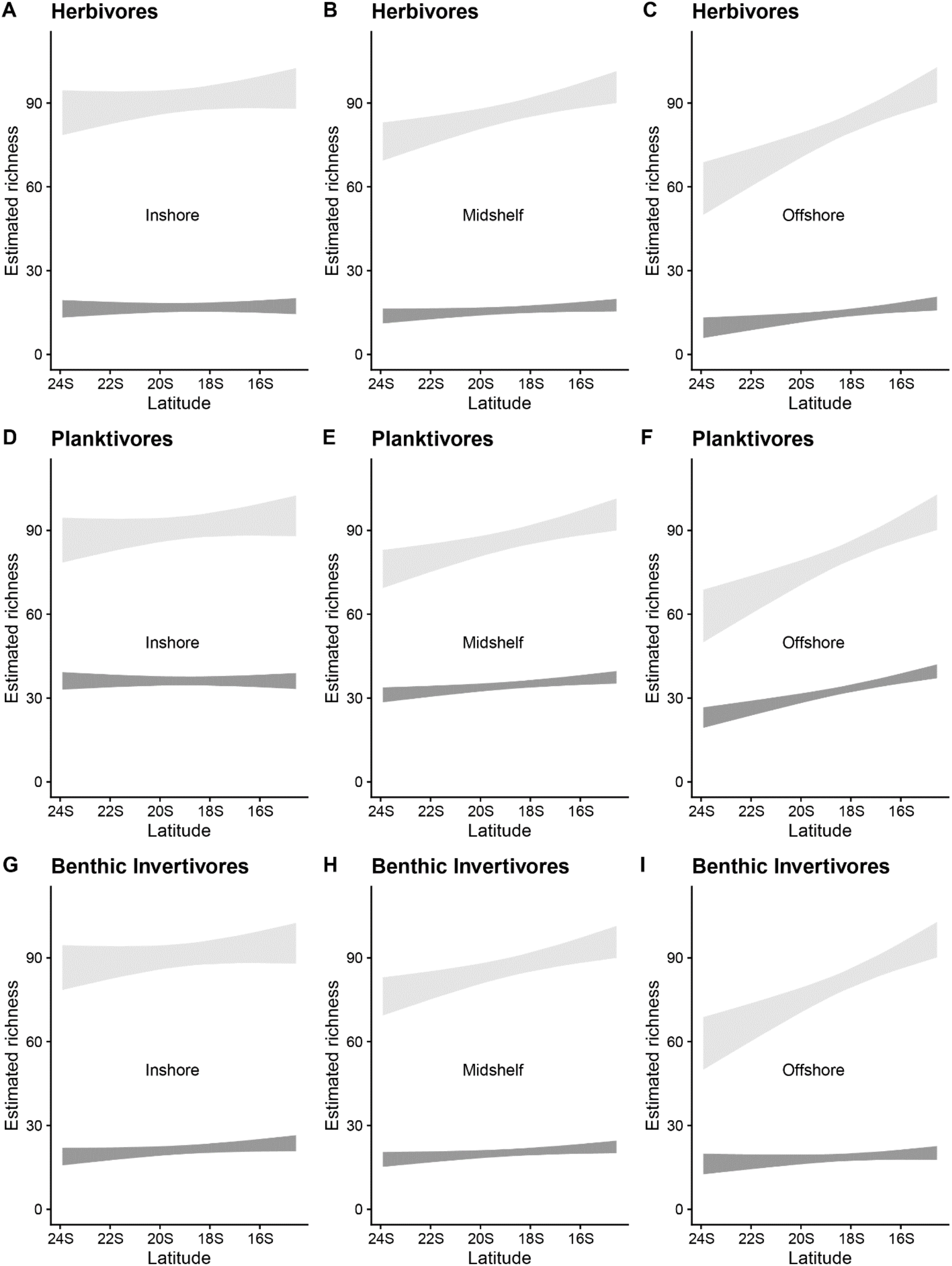
Richness of reef fish functional groups depend on latitude and cross-shelf position. Relationship between time-averaged species richness of reef fish trophic groups and the interaction of latitude with cross-shelf position. (**A**-**C**) Herbivores. (**D**-**F**) Planktivores. (**G**-**I**) Benthic invertivores. Grey bands are the 95% confidence intervals predicted from the lowest-AIC models for richness (n=40 reefs; Table S6). To better illustrate the interactive relationships, the 1^st^, median, and 3^rd^ quartiles of cross-shelf positions are fixed in panels (**A**-**C**), (**D**-**F**), and (**G**-**I**), respectively, and the relationship between species richness as a function of latitude show for the corresponding value of cross-shelf position. Note that cross-shelf position increases towards the coast.

**Fig. S7.**
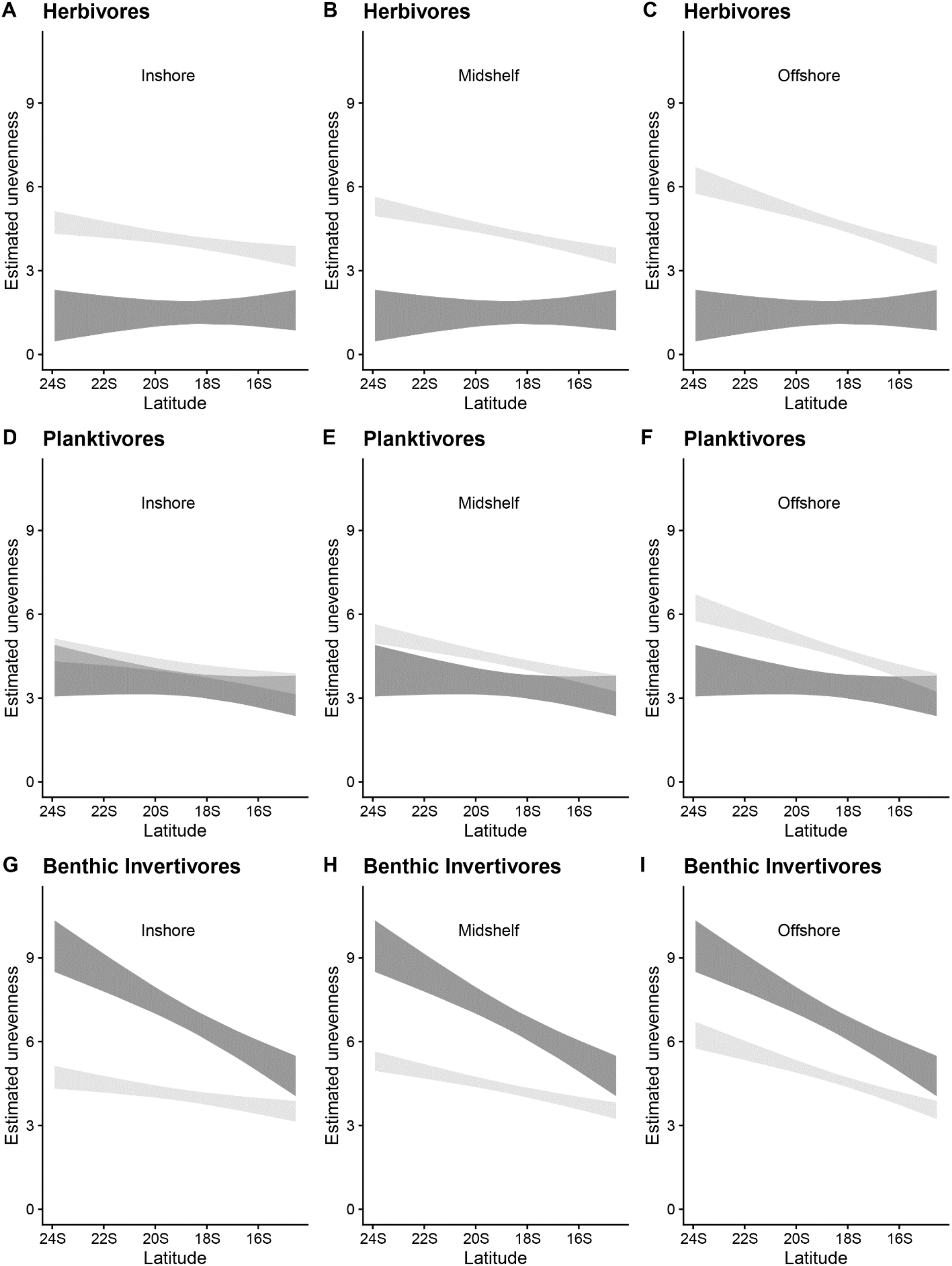
Unevenness of reef fish functional groups depend on latitude and cross-shelf position. Relationship between time-averaged unevenness of reef fish trophic groups and the interaction of latitude with cross-shelf position. (**A**-**C**) Herbivores. (**D**-**F**) Planktivores. (**G**-**I**) Benthic invertivores. Grey bands are the 95% confidence intervals predicted from the lowest-AIC models for unevenness (n=40 reefs; Table S6). To better illustrate the interactive relationships, the 1^st^, median, and 3^rd^ quartiles of cross-shelf positions are fixed in panels (**A**-**C**), (**D**-**F**), and (**G**-**I**), respectively, and the relationship between unevenness as a function of latitude show for the corresponding value of cross-shelf position. Note that cross-shelf position increases towards the coast.

**Fig. S8.**
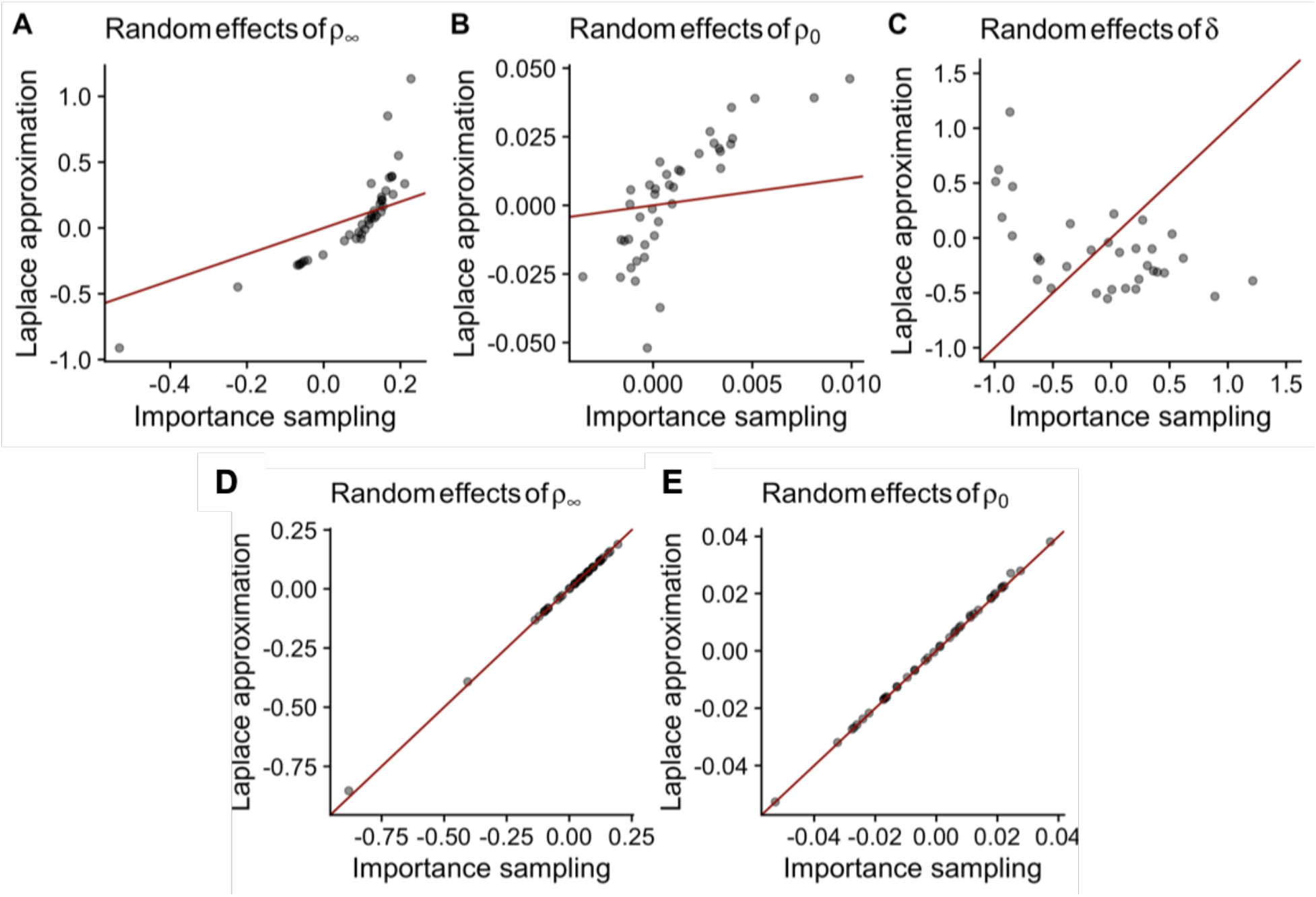
Relationship between random-effect parameters estimated from Laplace approximation and importance sampling in the full random effects model vs. the lowest-AIC random effects model. In the full model (**A**-**C**), density dependence *δ* is assumed to be a random-effect parameter in addition to the other two parameters of eq. 1 in the main text. By contrast, in the best model (**D**-**E**), density dependence *δ* is a fixed effect while keeping the other two parameters of eq. 1 as random effects. Red line represents a 1:1 relationship where estimates from both methods of Laplace approximation and importance sampling converge to the same values. The strong departures from the red line in panels (**A**-**C**) indicate numerical instability of the estimated values for this model and illustrate why we excluded this model from our model selection procedure (Table S10).

### Supplementary Tables S1 to S10

**Table S1.** OLS regression models for proportional variance components, richness, and unevenness. For explanatory variables, ‘mC’ and ‘log(sdC)’ represents the long-term (11-yr) mean (average coral cover) and log-transformed standard deviation in annual coral cover fluctuations (coral cover volatility), respectively. ‘Lat’ and ‘Shelf’ represents latitude and cross-shelf position, respectively. ‘Intercept’ represents the regression model that contains only an intercept. Cross symbols indicate models that include main effects and interactions, whereas plus symbols denote models including only main effects (i.e., additive effects of the explanatory variables). The analyses of the variance component of demographic and sampling variance are not presented, because their magnitudes are negligible compared to those of other variance components (Fig. 1E in the main text).

| Response variable | Explanatory variable | adj$R^2$ | AIC | ΔAIC |
| --- | --- | --- | --- | --- |
| Persistent species differences | log(sdC)×mC | 0.39 | -80.93 | 0 |
|  | log(sdC)×mC+Lat×Shelf | 0.34 | -75.66 | 5.27 |
|  | log(sdC) | 0.19 | -71.64 | 9.29 |
|  | mC | 0.16 | -70.28 | 10.65 |
|  | log(sdC)+Lat×Shelf | 0.19 | -68.83 | 12.1 |
|  | mC+Lat×Shelf | 0.12 | -65.59 | 15.34 |
|  | Intercept | 0 | -64.2 | 16.73 |
|  | Shelf | 0 | -63.78 | 17.15 |
|  | Lat | 0 | -62.24 | 18.69 |
|  | Lat×Shelf | 0 | -59.87 | 21.06 |
| Environmental stochasticity | log(sdC)×mC | 0.4 | -77.47 | 0 |
|  | log(sdC)×mC+Lat×Shelf | 0.35 | -71.96 | 5.51 |
|  | log(sdC) | 0.19 | -67.43 | 10.04 |
|  | mC | 0.16 | -65.91 | 11.56 |
|  | log(sdC)+Lat×Shelf | 0.18 | -64.39 | 13.08 |
|  | mC+Lat×Shelf | 0.12 | -61.05 | 16.42 |
|  | Intercept | 0 | -59.85 | 17.62 |
|  | Shelf | 0 | -59.4 | 18.07 |
|  | Lat | 0 | -57.97 | 19.5 |
|  | Lat×Shelf | 0 | -55.55 | 21.92 |
| Richness | Lat×Shelf | 0.55 | 299.34 | 0 |
|  | mC+Lat×Shelf | 0.55 | 299.83 | 0.49 |
|  | log(sdC)+Lat×Shelf | 0.54 | 301.18 | 1.84 |
|  | log(sdC)×mC+Lat×Shelf | 0.54 | 303.61 | 4.27 |
|  | Shelf | 0.24 | 318.69 | 19.35 |
|  | Lat | 0.1 | 325.51 | 26.17 |
|  | Intercept | 0 | 328.96 | 29.62 |
|  | mC | 0.01 | 329.39 | 30.05 |
|  | log(sdC) | 0 | 330.47 | 31.13 |
|  | log(sdC)×mC | 0 | 332.25 | 32.91 |
| Unevenness | mC+Lat×Shelf | 0.7 | 60.45 | 0 |
|  | log(sdC)+Lat×Shelf | 0.7 | 60.58 | 0.13 |
|  | Lat×Shelf | 0.71 | 60.78 | 0.33 |
|  | log(sdC)×mC+Lat×Shelf | 0.69 | 63.8 | 3.35 |
|  | Lat | 0.27 | 94.14 | 33.69 |
|  | Shelf | 0.22 | 96.93 | 36.48 |
|  | log(sdC) | 0.03 | 105.65 | 45.2 |
|  | Intercept | 0 | 106.03 | 45.58 |
|  | mC | 0.02 | 106.16 | 45.71 |
|  | log(sdC)×mC | 0.02 | 107.83 | 47.38 |

**Table S2.**
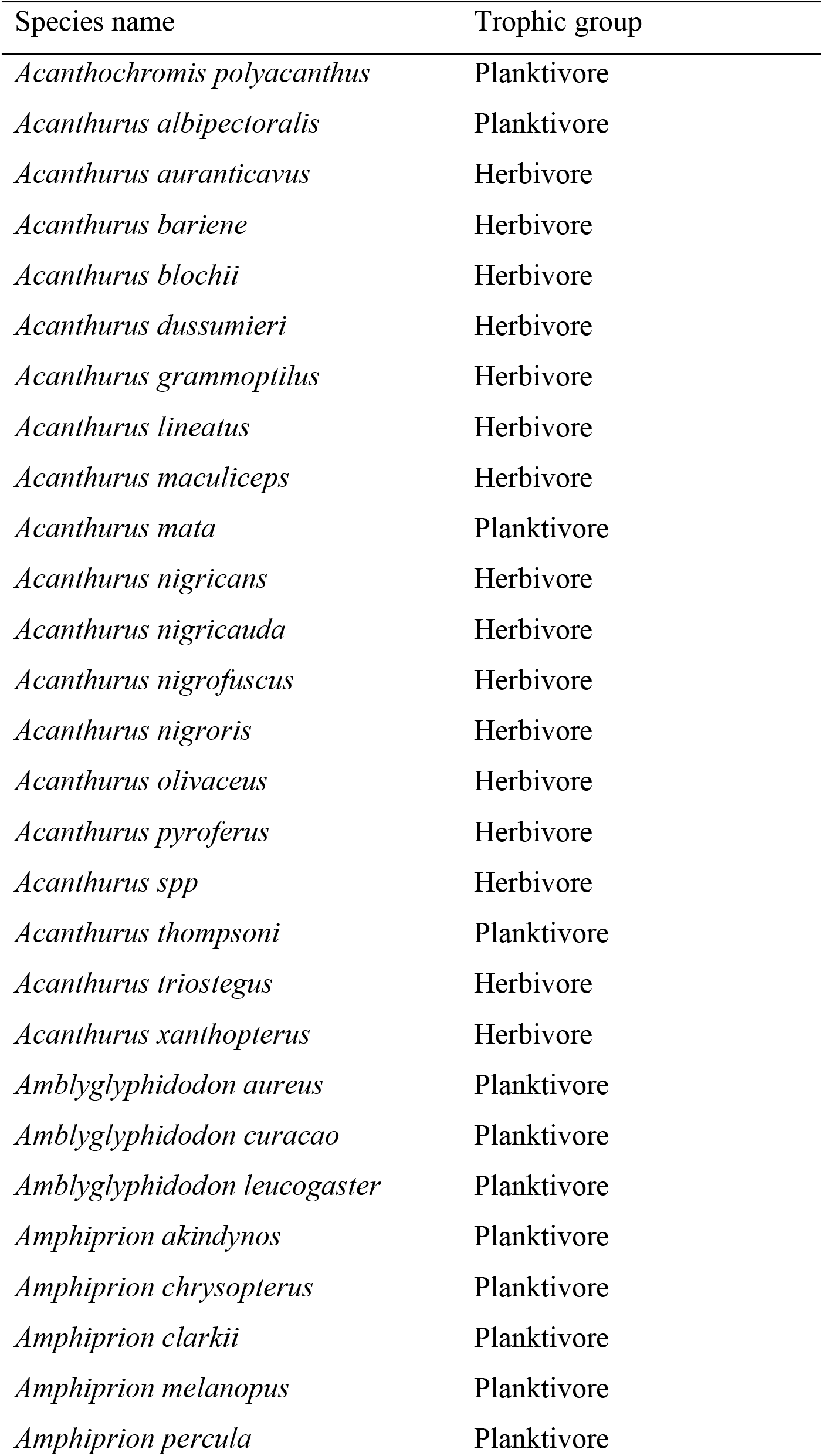
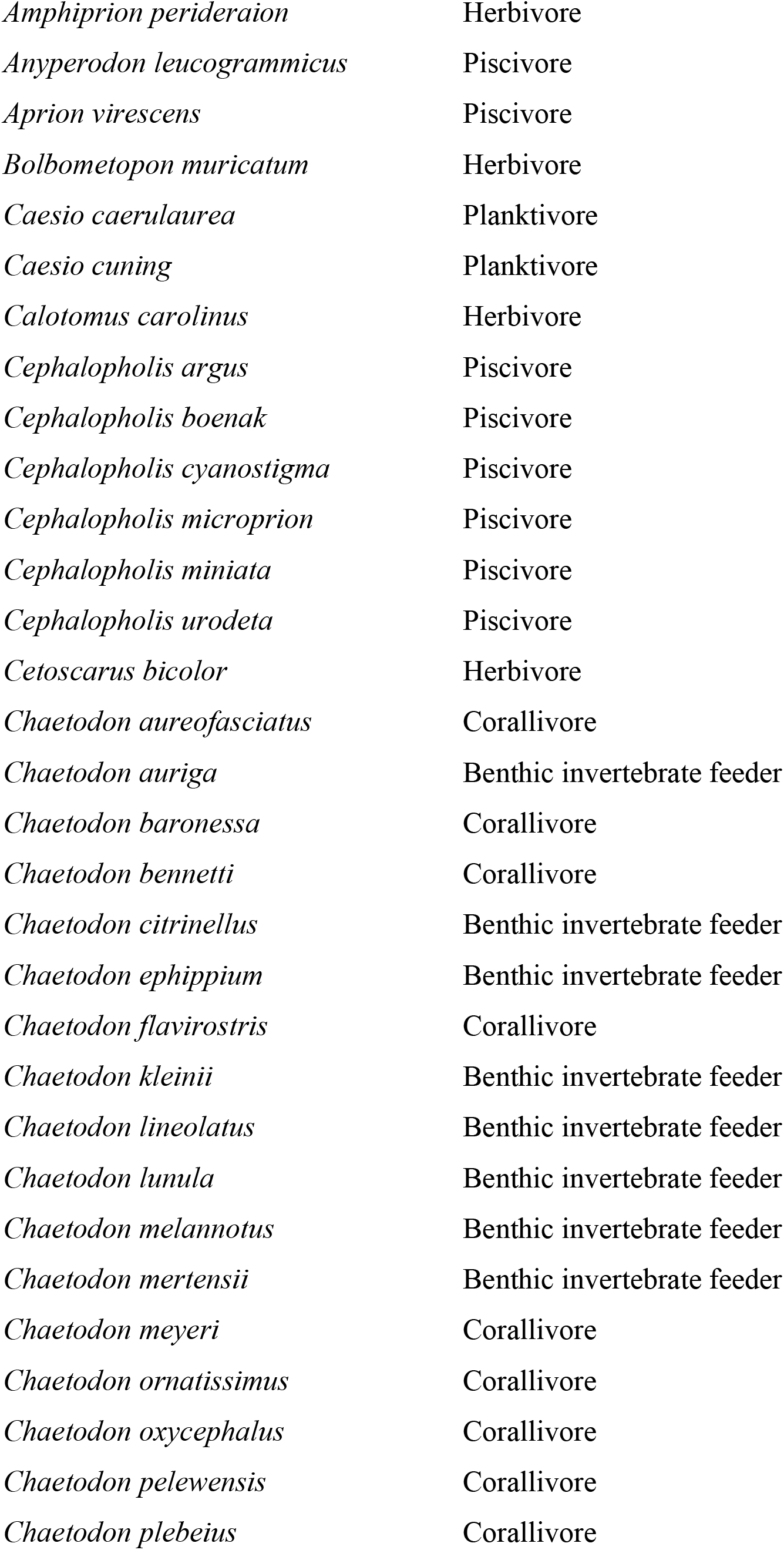
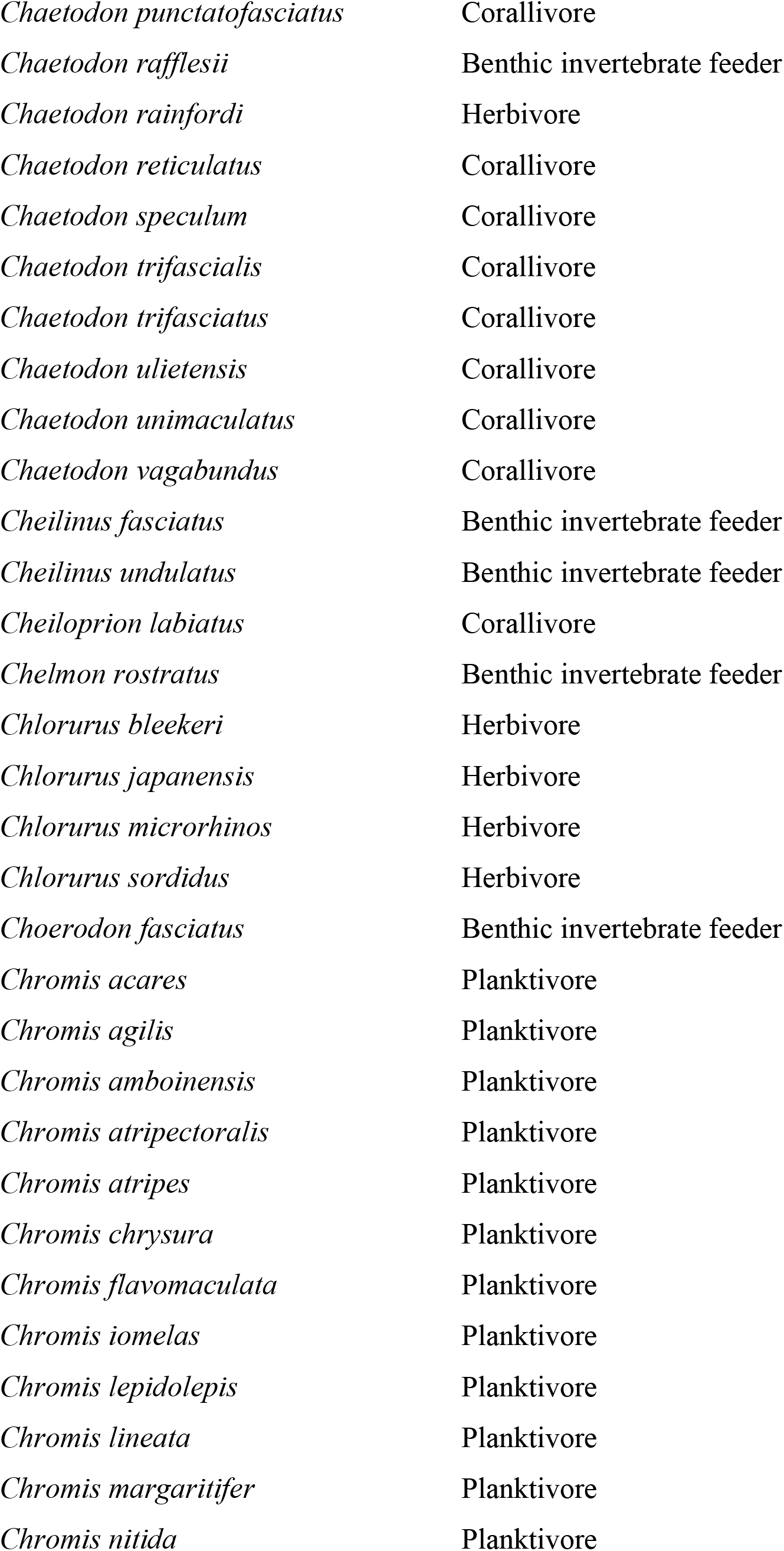
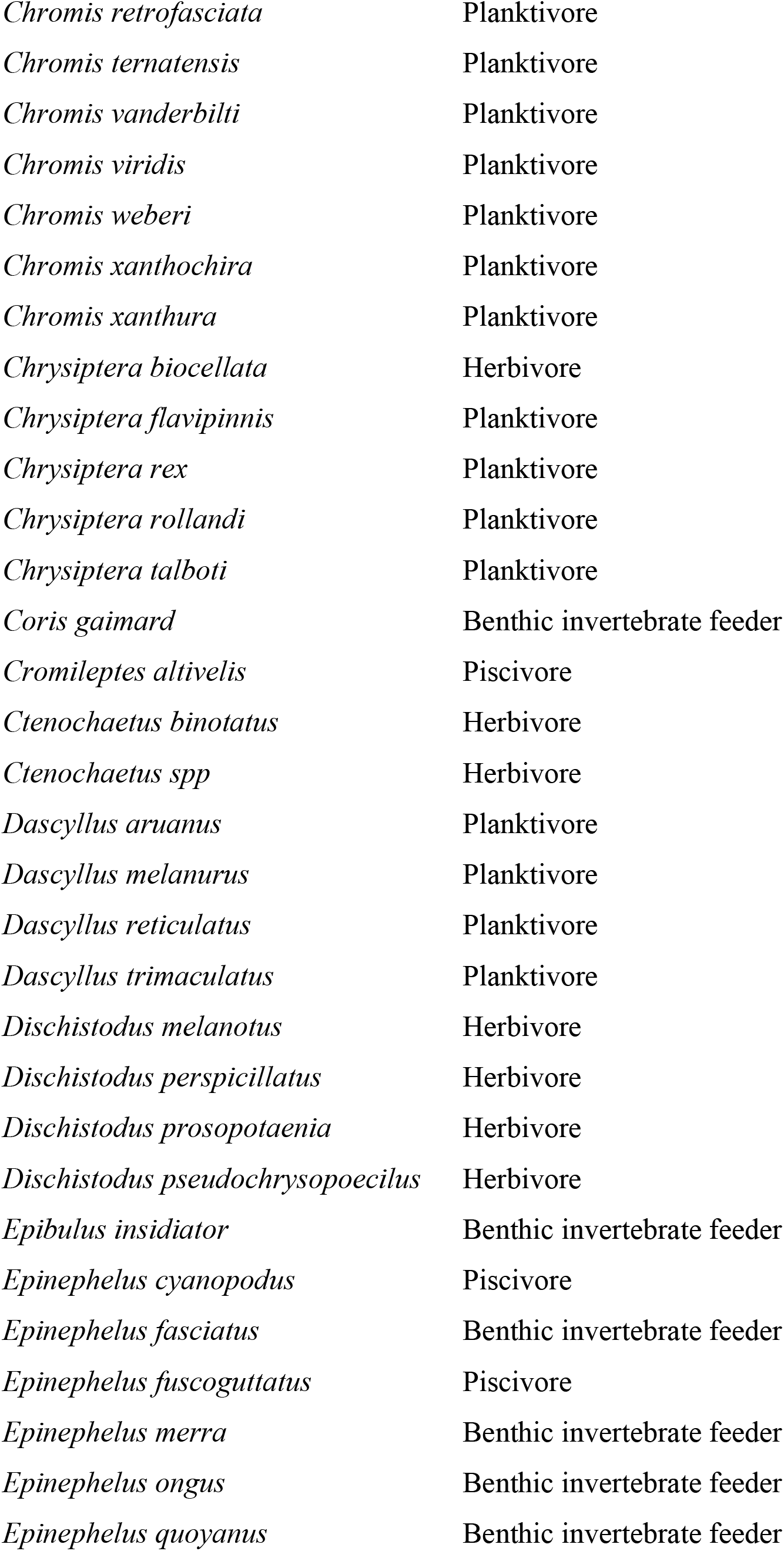
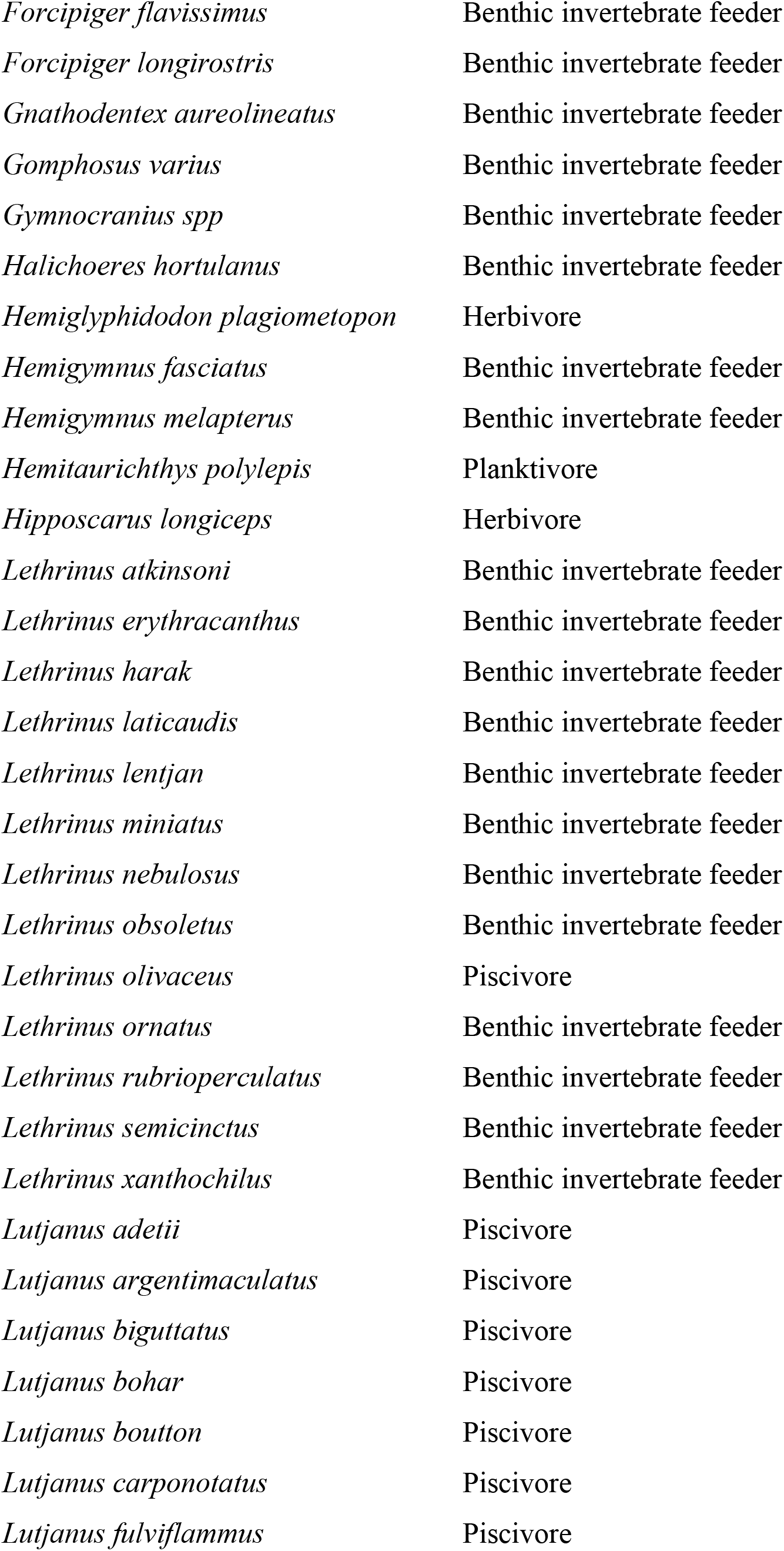
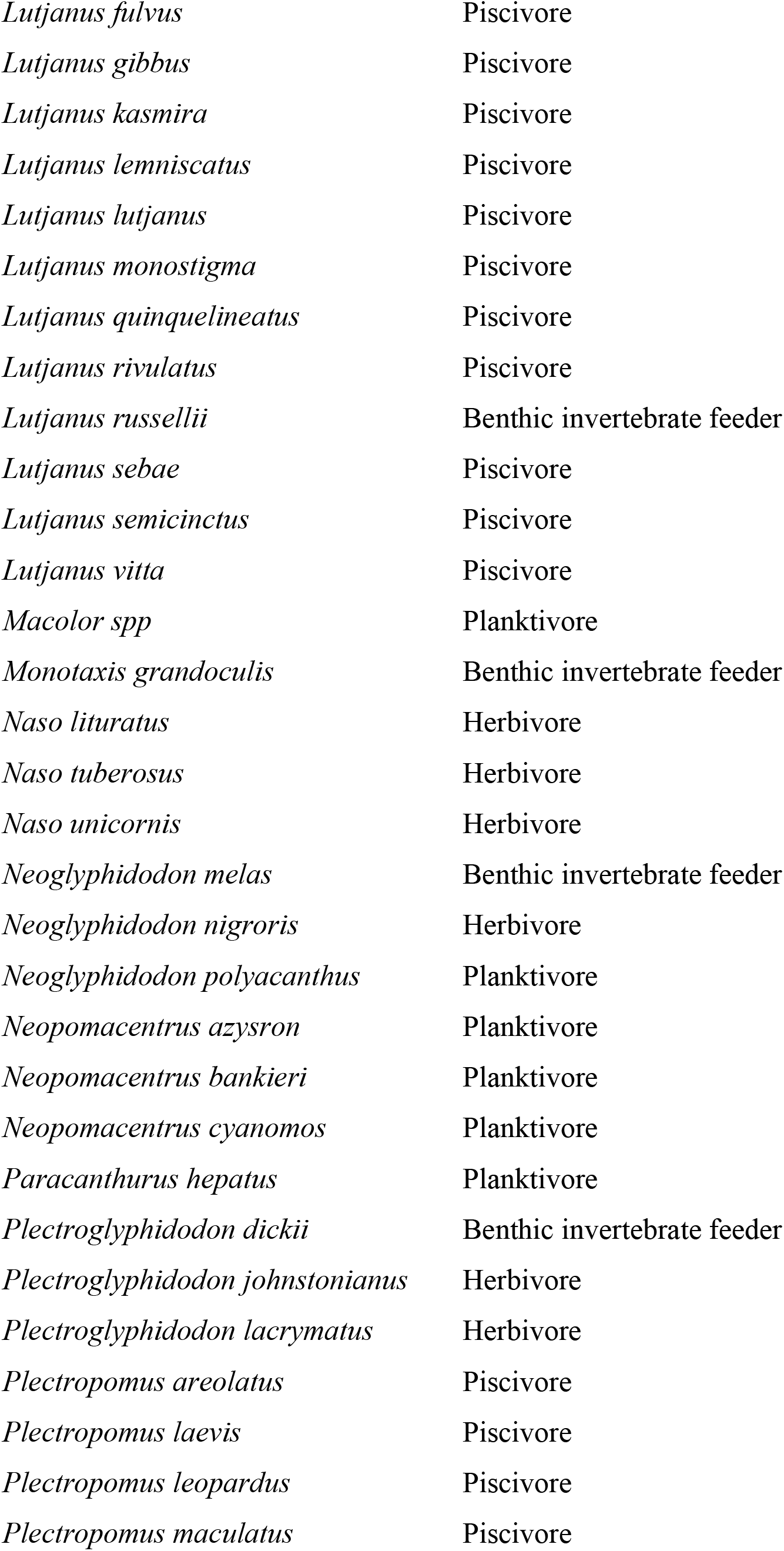
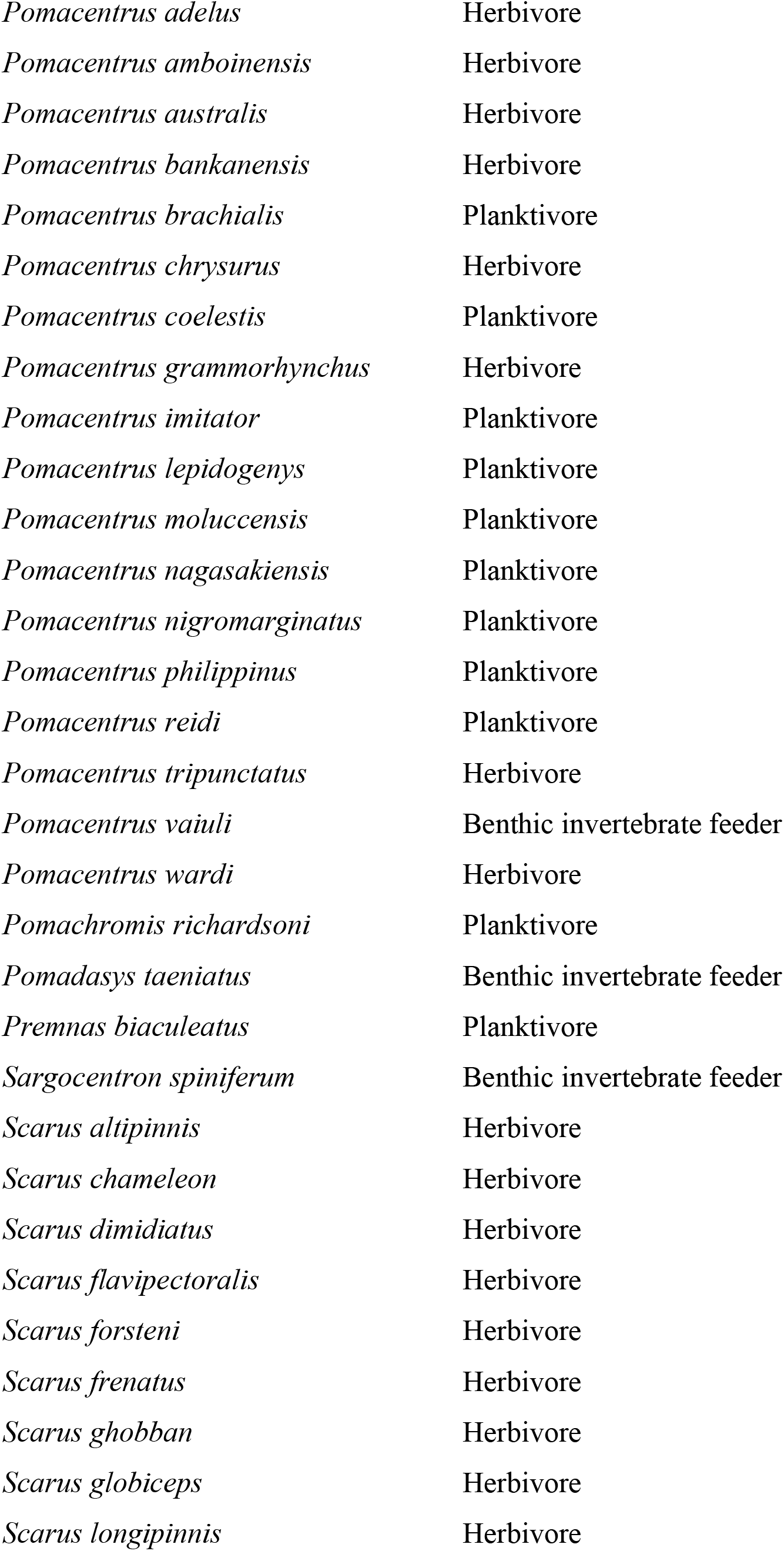
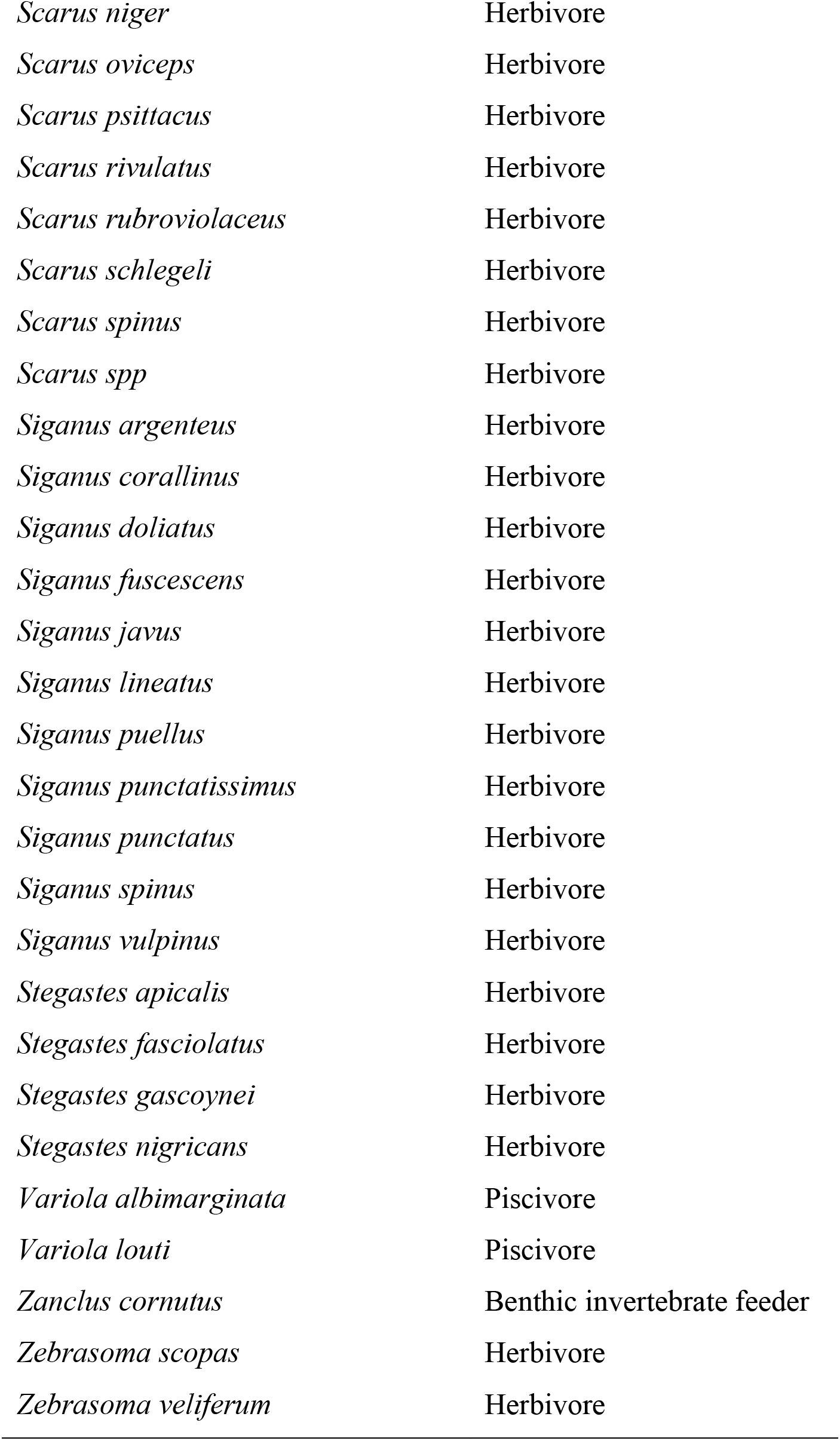
Classifications of trophic/functional groups of coral reef fishes on the Great Barrier Reef.

**Table S3.** Results of paired t-tests for differences between variance components of intrinsic species differences of trophic groups. The direction of the estimated mean difference is presented as row subtracted by column. Thus, for example, −1.8%, in the herbivore row and plantkivore column, indicates that the species differences explained 1.8% less variation for herbivores than it did for planktivores. * indicates *P* <0.05 and ** indicates *P* <0.01 after Bonferroni correction of *P* values.

|  | Herbivore | Planktivore | Benthicinvertivore |
| --- | --- | --- | --- |
| Herbivore | NA | -1.8% | 1.9% |
| Planktivore | NA | NA | 3.7% |
| Benthicinvertivore | NA | NA | NA |

**Table S4.**
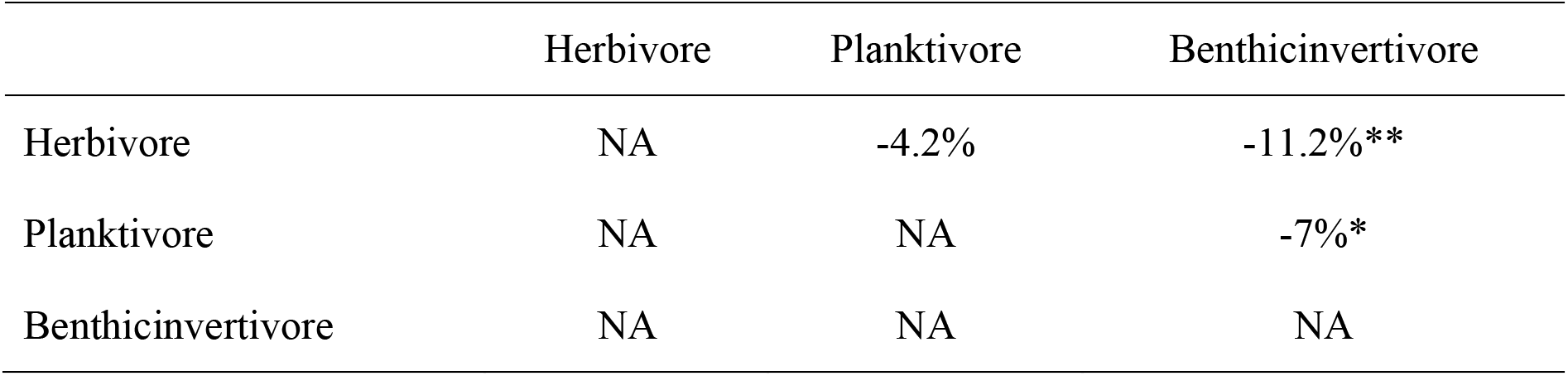
Results of paired *t*-test for differences between variance components of environmental stochasticity of trophic groups. The direction of the estimated mean difference is presented as per Supplementary Table S3. * indicates *P* <0.05 and ** indicates *P* <0.01 after Bonferroni correction of *P* values.

**Table S5.** Results of paired t-test for difference between variance components of overdispersion of trophic groups. The direction of the estimated mean difference is presented as per Supplementary Table S3. * indicates *P* <0.05 and ** indicates *P* <0.01 after Bonferroni correction of *P* values.

|  | Herbivore | Planktivore | Benthicinvertivore |
| --- | --- | --- | --- |
| Herbivore | NA | 6.1%** | 8.8%** |
| Planktivore | NA | NA | 2.6%** |
| Benthicinvertivore | NA | NA | NA |

**Table S6.**
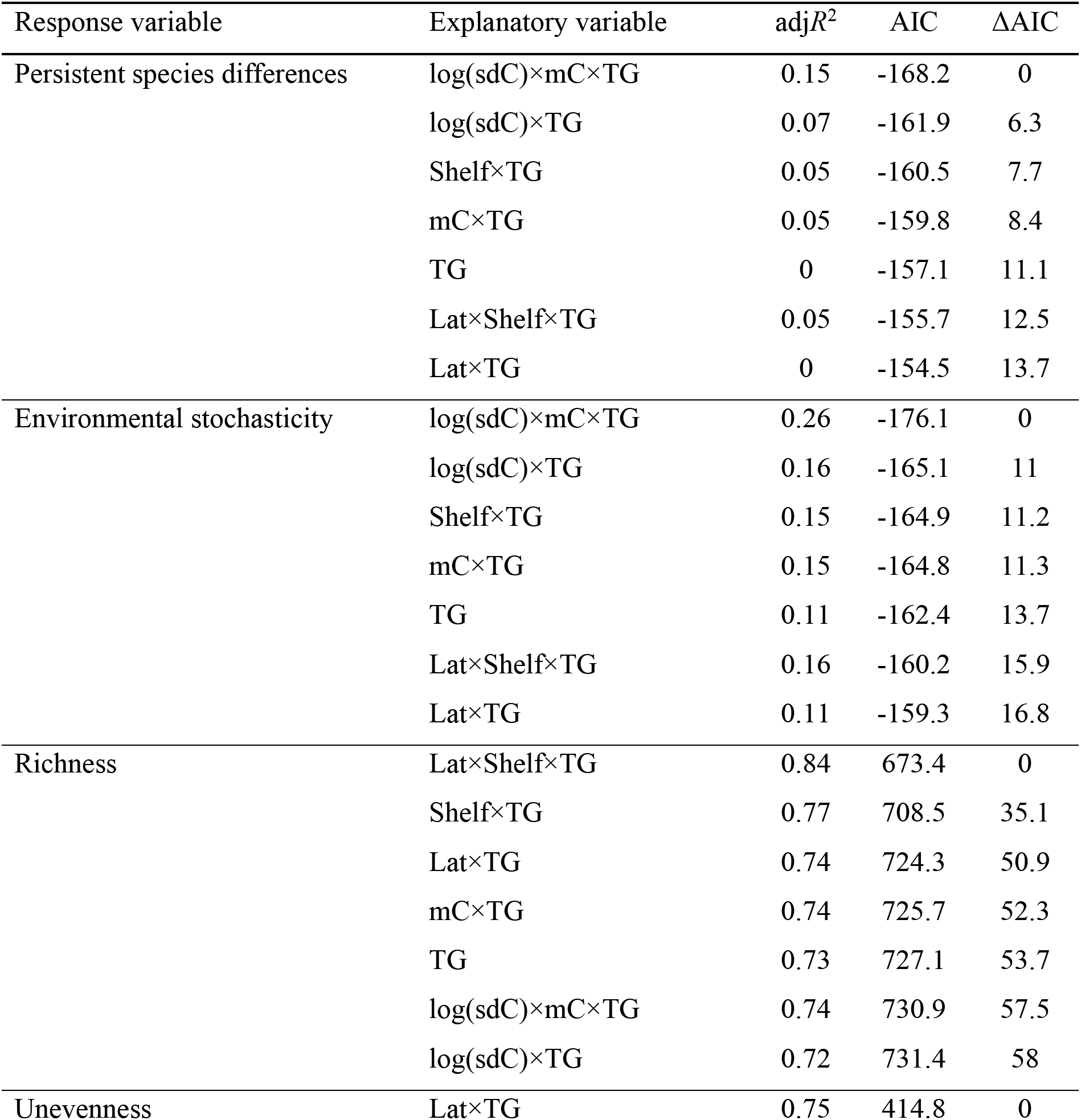
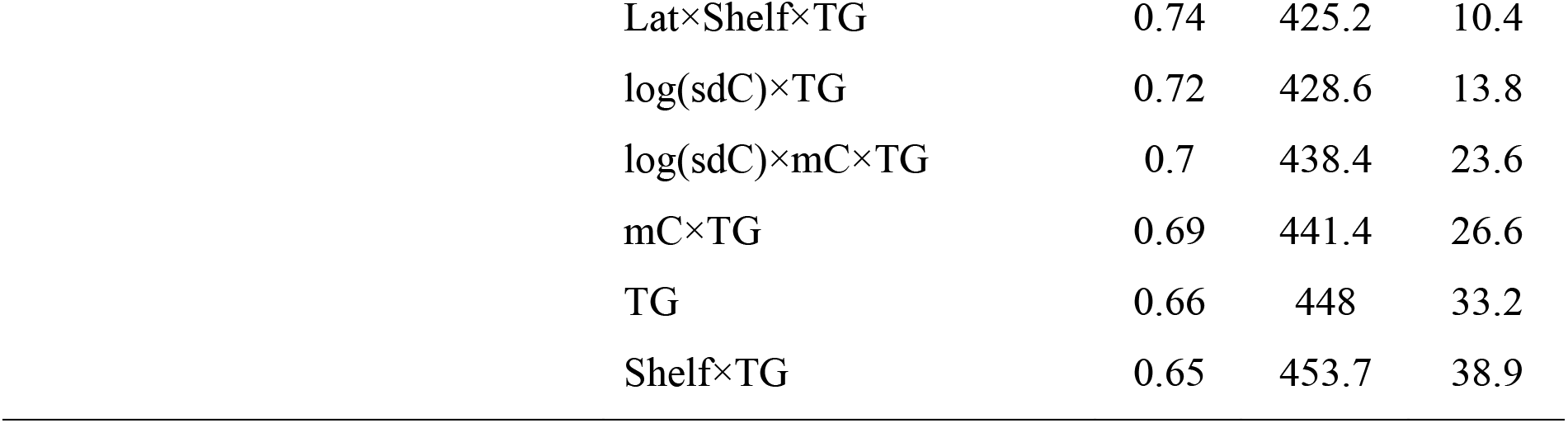
OLS regression models for proportional variance components, richness, and unevenness in the context of trophic groups. For explanatory variables, ‘mC’ and ‘log(sdC)’ represents the long-term (11-yr) mean and log-transformed standard deviation in annual coral cover fluctuations, respectively. ‘Lat’ and ‘Shelf’ represents latitude and cross-shelf position, respectively. ‘TG’ represents the identity of trophic group as a categorical variable. Cross and plus symbols represent interactive and additive effects, respectively. The analyses of the variance component of demographic and sampling variance (overdispersion) are not presented, because their magnitude is negligible compared to that of other variance components (Fig. S3).

| Response variable | Explanatory variable | adj$R^2$ | AIC | ΔAIC |
| --- | --- | --- | --- | --- |
| Persistent species differences | log(sdC)×mC×TG | 0.15 | -168.2 | 0 |
|  | log(sdC)×TG | 0.07 | -161.9 | 6.3 |
|  | Shelf×TG | 0.05 | -160.5 | 7.7 |
|  | mC×TG | 0.05 | -159.8 | 8.4 |
|  | TG | 0 | -157.1 | 11.1 |
|  | Lat×Shelf×TG | 0.05 | -155.7 | 12.5 |
|  | Lat×TG | 0 | -154.5 | 13.7 |
| Environmental stochasticity | log(sdC)×mC×TG | 0.26 | -176.1 | 0 |
|  | log(sdC)×TG | 0.16 | -165.1 | 11 |
|  | Shelf×TG | 0.15 | -164.9 | 11.2 |
|  | mC×TG | 0.15 | -164.8 | 11.3 |
|  | TG | 0.11 | -162.4 | 13.7 |
|  | Lat×Shelf×TG | 0.16 | -160.2 | 15.9 |
|  | Lat×TG | 0.11 | -159.3 | 16.8 |
| Richness | Lat×Shelf×TG | 0.84 | 673.4 | 0 |
|  | Shelf×TG | 0.77 | 708.5 | 35.1 |
|  | Lat×TG | 0.74 | 724.3 | 50.9 |
|  | mC×TG | 0.74 | 725.7 | 52.3 |
|  | TG | 0.73 | 727.1 | 53.7 |
|  | log(sdC)×mC×TG | 0.74 | 730.9 | 57.5 |
|  | log(sdC)×TG | 0.72 | 731.4 | 58 |
| Unevenness | Lat×TG | 0.75 | 414.8 | 0 |

|  |  |  |  |
| --- | --- | --- | --- |
| Lat×Shelf×TG | 0.74 | 425.2 | 10.4 |
| log(sdC)×TG | 0.72 | 428.6 | 13.8 |
| log(sdC)×mC×TG | 0.7 | 438.4 | 23.6 |
| mC×TG | 0.69 | 441.4 | 26.6 |
| TG | 0.66 | 448 | 33.2 |
| Shelf×TG | 0.65 | 453.7 | 38.9 |

**Table S7.** Pearson correlations between environmental variables. ‘mC’, ‘log(sdC)’, and ‘log(cvC)’ represents the long-term (11-yr) average, log-transformed standard deviation, and log-transformed coefficient of variation in coral cover annual fluctuations, respectively. ‘Lat’ and ‘Shelf’ represents the latitude and cross-shelf position, respectively. Symbol * indicates *P*<0.05.

|  | log(sdC) | mC | log(cvC) | Lat | Shelf |
| --- | --- | --- | --- | --- | --- |
| log(sdC) |  | 0.53 | 0.89* | -0.24 | -0.04 |
| mC |  |  | 0.12 | -0.16 | -0.12 |
| log(cvC) |  |  |  | -0.18 | 0.07 |
| Lat |  |  |  |  | 0.15 |
| Shelf |  |  |  |  |  |

**Table S8.**
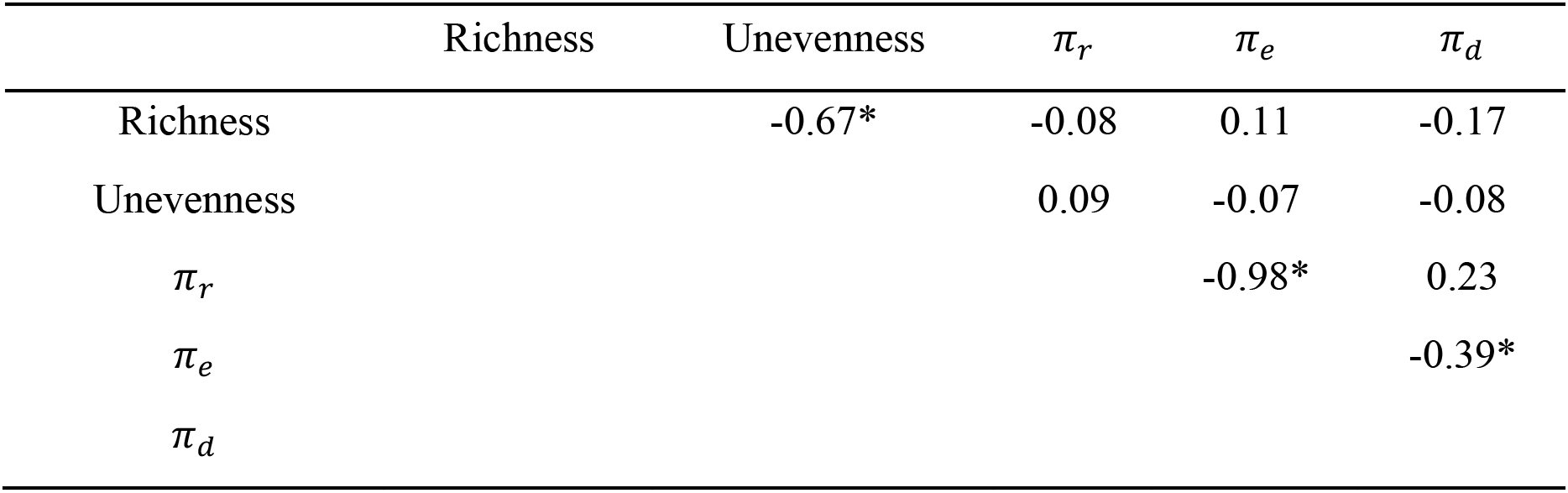
Pearson correlations between static and dynamic measures of reef fish community structure. ‘Richness’ and ‘Unevenness’ represents time-averaged richness and unevenness from Poisson-lognormal fits of relative species abundance of reef fish across coral reefs, respectively. *π_r_*, *π_e_* and *π_d_* (as eqs 2-4 in the main text) represents the proportional variance in relative abundances of reef fishes explained by intrinsic species differences, environmental stochasticity, and demographic and sampling variance (overdispersion), respectively. Symbol * indicates *P*<0.05.

**Table S9.** Effect size comparison for AIC-similar (ΔAIC<2) models of unevenness. Variables are standardized prior to estimating effect size. Model structure for predicting unevenness presented as per Table S1. ** indicates *P* <0.01.

| Model | Intercept | Lat | Shelf | Lat×Shelf | mC | log(sdC) |
| --- | --- | --- | --- | --- | --- | --- |
| Unevenness~Lat×Shelf | 0.05 | -0.73** | 0.59** | -0.38** | NA | NA |
| Unevenness~Lat×Shelf+mC | 0.05 | -0.71** | 0.6** | -0.34** | 0.13 | NA |
| Unevenness~Lat×Shelf+log(sdC) | 0.05 | -0.71** | 0.59** | -0.4** | NA | 0.12 |

**Supplementary Table S10.** Model selection of auto-correlation function eq. 1 for VPRSA. *ρ*_∞_, *ρ*_0_, and δ indicate either the fixed-effect parameter estimate (where the corresponding “SD(.)” column is NA), or the estimated mean of the random-effect distribution, for the parameters of the autocorrelation function eq. 1 in the main text, respectively. SD(.) is the estimated standard deviations of the random-effect distribution for the corresponding parameter. SDtotal is the estimated residual standard error. 95% confidence intervals are enclosed in square brackets. Random effects are modelled as following lognormal distributions because normally distributed random effects yielded numerally unstable fits. Note that, with respect to all possible combinations of fixed and random-effect model structure of eq. 1., only the results of numerically stable models are reported. ‘AIC’ represents Akaike information criteria. Bootstrapped marginal-AIC values are consistent with AIC (not shown for simplicity).

| Fixed effect | Random effect | $\rho_\infty$ | $\rho_0$ | $\delta$ | SD ($\rho_\infty$) | SD ($\rho_0$) | SD<sub>total</sub> | AIC |
| --- | --- | --- | --- | --- | --- | --- | --- | --- |
| $\rho_\infty, \rho_0, \delta$ | $\rho_\infty, \rho_0$ | 0.75 [0.72, 0.78] | 0.93 [0.92, 0.94] | 0.09 [0.08, 0.11] | 0.18 [0.15, 0.22] | 0.02 [0.02, 0.03] | 0.05 [0.047, 0.051] | -6844.62 |
| $\rho_\infty, \rho_0, \delta$ | $\rho_\infty$ | 0.75 [0.73, 0.78] | 0.93 [0.93, 0.94] | 0.09 [0.08, 0.11] | 0.16 [0.13, 0.2] | NA | 0.05 [0.049, 0.051] | -6803.06 |
| $\rho_\infty, \rho_0, \delta$ | NA | 0.76 [0.7, 0.82] | 0.93 [0.93, 0.94] | 0.09 [0.05, 0.15] | NA | NA | 0.06 [0.059, 0.061] | -6096.7 |

### Supplementary Simulation Study

Our analyses of the reef fish data use the variance partitioning of relative species abundance (hereafter, VPRSA) method. However, the underlying community dynamics model from which VPRSA is derived (eq. 5-6 in the Methods) makes some important simplifying assumptions that may commonly be violated in real communities, including the data analyzed in this paper, so it is important to evaluate whether inferences about proportional variance components are sensitive to those assumptions. Specifically, we wish to determine whether estimates produced by the VPRSA method, when applied to simulated data that violate model assumptions, still produce variance component estimates that reflect the relative contribution of persistent intrinsic species differences versus stochastic fluctuations to overall variation in species abundances.

Simplifying assumptions of this model include that the following: (i) species’ intrinsic growth rates vary according to a normal distribution, (ii) the strength of intra-specific density dependence is the same for all species and inter-specific interactions to be negligible, and (iii) the responses of species’ intrinsic growth rates to environmental stochasticity (environmental variance) are assumed to be independent and equal in magnitude (i.e., they fluctuate from year to year with the same variance; because of the log-scaling of abundance in this model, this implies that fluctuations in population growth have a variance that is proportional to the mean). Samples from communities whose abundances follow these assumptions generate a static species-abundance pattern that follows a Poisson-lognormal distribution, consistent with what is commonly observed in data (*33, 42*), and allow derivation of the autocorrelation function that we used in the main text to partition the variance in relative species log-abundances into components.

Some of these assumptions are likely to be more reasonable than others. For instance, the assumption of Gompertz-type density dependence is consistent with many previous studies, which have found that this model characterizes the functional form of density dependence well and performs better than, or as well as, other forms such as the Ricker or Logistic form (*43–45*). More specifically, in a previous study of reef fish functional group dynamics on the Great Barrier Reef (3*5*), Gompertz-type density dependence was found to fit data better than other forms of density dependence. Another assumption is that of a normal distribution of intrinsic growth rates. Because of the log-scaling of species abundances in eq. 5, this implies a lognormal distribution of geometric growth factors. A strongly right-skewed distribution of this quantity, such as a lognormal, is consistent with the few studies of variation in population growth at the assemblage-level, which show that most species are relatively slow growing, with a long tail of a few fast-growing species (*46*).

In contrast, the assumptions of equal strength of density dependence, and equal proportional magnitude of environmentally induced fluctuations in abundance, seem unlikely to hold in nature (*35, 47*). Between these extremes, the assumptions that interspecific interactions are negligible, and that species respond independently to environmental fluctuations, are common in biodiversity models, but controversial. For instance, there is some evidence that between-species interactions tend to be weak, particularly for high-diversity systems (*35, 48, 49*), and species’ responses to fluctuations tend to be relatively independent on average (*32, 50*). As noted above, the lognormal shape of the static species abundance distribution has been shown previously to be robust to violation of these assumptions (*33, 51*), but whether variance components estimated from the temporal evolution of such species-abundance distributions are equally robust is unknown. For instance, the deterministic “persistent species difference” component captures the proportional variance in log-abundance due to species differences in equilibrium abundance, but this will no longer be directly proportional to variance in intrinsic growth rates (as in the *Vr* term of eq. 7 in the main text) when species interactions or among-species heterogeneity in density dependence is present.

In this supplement, we simulate different scenarios of community dynamics to test the robustness of variance components estimated from VPRSA to violations of assumptions. Specifically, we conduct VPRSA analysis on simulated community dynamics data that systematically violate model assumptions of VPRSA, and we compare estimated variance components with approximate “true” variance components based on the analytical solutions and known underlying parameters of the simulated communities. R code for simulations and fits is available at https://github.com/TsaiCH/simsEngenVPRSA. Our objective here is not to comprehensively examine the statistical performance of the estimates from this method, but rather to verify that any biases in the estimates produced when the model’s assumptions were violated do not compromise the conclusions drawn about the reef fish data (LTMP) in the main text. For that reason, we focus on simulated data that share key features of the LTMP data, with respect to species richness, number and length of replicate time series, and parameter values.

#### Community dynamics model

We use state-space models that incorporate different assumptions about community dynamics to produce simulated data, which we then analyze using the VPRSA approach applied in the main text. This allows us to evaluate the robustness of VPRSA estimates to violation of the assumptions of the community dynamics model from which it was derived. The R code for simulations of community dynamics, and VPRSA estimation, is available and open access at https://github.com/TsaiCH/simsEngenVPRSA.

First, we simulate the abundance dynamics of species *i*=1..*S* in a community according to the discrete-time multivariate Gompertz model (*52*):

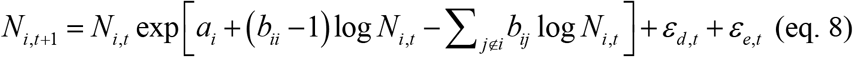

where *N_i_*_,*t*_ is the abundance of species we at time *t*, *a_i_* is the species-specific intrinsic growth rate, and *b_ii_* and *b_ij_* are coefficients related to intra- and inter-species density dependence, respectively. *b_ii_* −1 is the strength of intra-specific density-dependence, while larger values of *b_ij_* indicate stronger inter-specific interactions. Here, we constrain these values to be between 0 and 1, consistent with parameter estimates from Gompertz model fits to individual population time series (*16*), as well as the parameter estimates from our analyses here. (Note that *b_ii_* <0 represents strongly over-compensatory interactions, where increases in *Nt* produce decreases in *Nt*+1, whereas *bii*=1 indicates density-independent dynamics.) Additionally, the species-specific intrinsic growth rates ( *a_i_*) are assumed to vary among species according to a normal distribution (*sensu r_i_* in eqs.1-2). ɛ*_d,t_* and ɛ*_e,t_* are species-specific noise terms representing demographic and environmental stochasticity.

More formally, let log *N_t_* = **X_t_** and take the natural logarithm of both sides of eq. 8 in order to facilitate expressing the Gompertz-type community dynamics in matrix form. By doing so, the community dynamics model becomes an order-one multivariate autoregressive model with two components of process noise as follows:

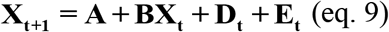

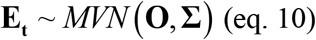

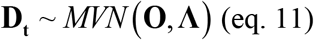

where **X_t_** is a vector containing log abundance for each species at time *t*, **A** is a vector containing species-specific intrinsic growth rates (*ai*), and **B** is the interaction matrix where the diagonal (*b_ii_*) and off-diagonals (*b_ij_*) are coefficients related to intra- and inter-specific density dependence (keeping in mind that the intra-specific density-dependence is *b_ii_* −1, whereas the intra-specific density dependence is *b_ij_*). **E_t_** is a vector of random variables containing species’ responses to environmental fluctuations (i.e., the perturbations to the intrinsic growth rate due to environmental stochasticity) and follows a multivariate normal distribution (*MVN*) with zero means and variance-covariance matrix **Σ** (eq. 10). **D_t_** is a vector of random variables representing perturbations due to demographic stochasticity. These quantities also follow a multivariate normal distribution (*MVN*) with zero means and variance matrix **Λ** (by definition, the covariances of this matrix are all zero). Because less abundant species are more prone to demographic stochasticity than abundant species, we follow previous work and model the demographic variances in log-abundance (the diagonal in **Λ**) as inversely proportional to the square root of species abundance (eq. 11) (*32, 53*). That is, the diagonal elements of **Λ**

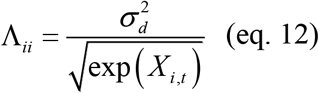

Finally, let **λ** represent a vector containing the expected relative species abundance in a random sample of the species-abundance distribution, such that:

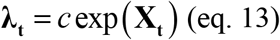

where *c* is a measure of sampling intensity. Hence the (simulated) sampled species abundances Yt will be a Poisson sample of **λ_t_**:

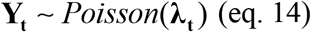

The final model of community dynamics represents a discrete-time multivariate state-space model with normally distributed equilibrium log-abundances, normally distributed process noise, and Poisson-distributed observation error. we therefore model the sampled abundance values *yt* as following a Poisson-lognormal distribution (*51*).

#### Simulating empirically constrained community dynamics data

Simulated data were constrained to have similar numbers of locations (“reefs”), time series lengths, species richness, and numbers of individuals as per the LTMP data analyzed in the main text. Specifically, we generated 100 simulated data sets, each of which consisted of 40 simulated time series (“reefs”) sampled annually for 11 years, to correspond to the time series for the 40 annually sampled reefs in the LTMP. For each reef, true total species richness was fixed at *S*=100 in all simulations. The level of “true” total richness used in simulations is close to the upper bound of estimated total richness at the reef scale in the LTMP (cf. Fig. 1 in the main text). Then, for each year at each reef, we simulated a Poisson random sample with a mean of 1500 individuals (i.e., the sampling intensity *c* was set so that the sum of the Poisson mean abundance across all species was equal to 1500), since this was close to the median sample size in the LTMP. If any simulated samples had fewer than 40 observed species (i.e., species with sampled abundance greater than zero), that sample was discarded, and a new random sample was taken from the community for that reef and year. This threshold of 40 observed species was used to prevent unrealistically low representation of the community (in the LTMP data, no reefs had fewer than 40 observed species in any year). Simulations where this occurred were extremely rare (approximately 2% of simulations), so it is unlikely that this culling process has affected our conclusions.

For each simulated data set, the communities on all reefs were specified to have the same community dynamics parameters for eqs. 9-11, except for the strength of environmental stochasticity (i.e., *σ_e_* in eq. 1 and the diagonal 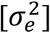 of **^Σ^** in eq. 10). This last quantity was varied systematically, in order to produce a data set in which the relative importance of deterministic species differences versus environmental stochasticity varied widely among reefs, which could then be used to evaluate how well VPRSA resolved these differences. Specifically, the environmental variance term (i.e., 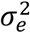) was varied from 0.025 to 0.5 in equal sized increments across the simulated reefs (e.g., one reef had 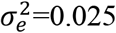, another had 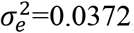, and so on up to 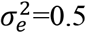). This created a true distribution of variance components among reefs that was uniform and extended almost all the way to zero. For the effect of persistent, intrinsic species differences (cf. eq. 1-2 and eq. 9), the species-specific intrinsic growth rates were modelled as varying among species according to a normal distribution with mean *μ_r_* =1.5 and standard deviation *σr* = 0.25, based on a meta-analysis of global fishery stock assessments (*46*). Demographic stochasticity was simulated as process noise (**D_t_**), where the demographic variances (the diagonal of **Λ** in eq. 11) were scaled by the value 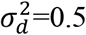. Other parameter values varied among simulation scenarios, as specified below.

#### Scenarios of simulated community dynamics data

We simulated five scenarios of community dynamics, which are constrained as described above, to test the robustness of variance components (i.e., the relative importance of deterministic versus stochastic factors in eq. 7 in the main text) estimated by VPRSA. The values of 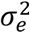 and 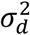 specified above were used for all simulations, and the additional parameters specified in scenarios (i)-(v) below were chosen so that the simulations produced frequency distributions of 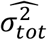 (variance of log-abundance in the communities), and sample completeness (measured as the fraction of the species pool observed at each site in each year) that were similar to those produced when the Poisson-lognormal was fitted to the LTMP data (Simulation Study Fig. 1). In addition, to ensure a stationary distribution of population sizes (i.e., all species coexisting), the complex norm of eigenvalues of interaction matrix **B** was constrained to be less than one. This constrained the overall strength of interspecific competition in scenario (iii) (i.e., if interactions were too strong, species would be unable to coexist).

The details of parameters and scenarios of simulated community data are as follows:

i. *Baseline*: These simulations were run to conform with the assumptions of the stochastic community-dynamic theory from which VPRSA was derived. Specifically, intra-specific density dependence was the same for all species (*bii*=*b*), the interaction matrix **B** contained no inter-specific density dependence (i.e., the off-diagonals of **B** were zero in eq. 9) and responses to environmental fluctuations were independent and equal in variance (i.e., the off-diagonals of **Σ** were zero in eq. 10, and the variances were all equal to the reef-specific values of 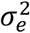 specified above: 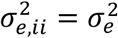). In these simulations, we set *b*=0.84 (i.e., the “strength of density dependence” was 1 - 0.84 = 0.16), as this is consistent with the estimate of this parameter from the real data.
ii. *Varied intra-specific interactions (varIntra)*: This scenario introduces between-species variation in intra-specific density dependence. Specifically, values on the diagonal of **B** in (eq. 9) were drawn from a normal distribution with mean 1.8 and standard deviation 0.4, and then inverse-logit transformed to yield values between 0 and 1. This produces random coefficients of the diagonal of **B** with mean values close to 0.84 (i.e., *E*[*b_ii_*] ≈ 0.84, implying average strength of density-dependence 1 − *E*[*b_ii_*] ≈0.16), and standard deviations close to 0.06, implying a coefficient of variation of density-dependent strength of about 0.37. Because equilibrium abundance is exp 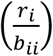, this approach produced unrealistically large variance in the total variance of log-abundance, 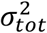. Therefore, the diagonal elements of **B** werereordered to increase with species’ intrinsic growth rates (i.e., elements of vector **A** in eq. 9), so that species with strong density dependence also had high intrinsic growth rates. This yielded more realistic variances of log-abundance (see below).
iii. *Inter-specific interactions (varInter)*: This scenario introduces diffuse inter-specific density dependence by drawing the off-diagonals of **B** from a uniform distribution between 0 and 0.002 (mean=0.001). This yielded an average summed effect of interspecific interactions (across the other 99 species in the community) that was approximately 60% the strength of intra-specific density-dependence (i.e., 0.001 × 99 ≈ 0.6 × 0.16). All other parameter values were the same as in the Baseline simulation. Mean interaction strength values slightly above those employed here (0.001-0.003) tended to produce distributions of observed richness values that differed notably from the data (lower observed richness levels, and more strongly right-skewed abundance distributions). Moreover, mean interaction strengths above about 0.003 tended to produce assemblages lacking a stable coexistence equilibrium.
iv. *Unequal environmental variances (varEnv)*: This scenario introduces heterogeneity among species in sensitivity to environmental fluctuations. Specifically, species-specific environmental variances (the diagonal of Σ in eq. 10) were drawn from a uniform distribution between 0 and 2 (mean=1), and then multiplied by the reef-specific 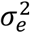 term, as specified above. This ensured that the average value of environmental variance ranged from 0.025 to 0.5, as in the *Baseline* scenario, and thus continued to yield realistic 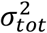 values. All other parameter values were as in the *Baseline* scenario.
v. *Unequal environmental covariances (varcovEnv)*: This scenario introduces covariance in species’ responses to environmental fluctuations. Specifically, we generated a lower-triangular matrix **L** whose elements were drawn from a normal distribution with mean 0 and standard deviation 0.25. We then produced a covariance matrix **Σ** = **LL^T^**. The elements of **Σ** were subsequently standardized by the mean of the diagonal elements, and then the entire matrix multiplied by the reef-specific environmental variance term 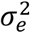. This yielded a matrix of unequal variances and covariances among species, whose correlation coefficients ranged from −0.5 to 0.5, with mean 0, and with the mean of the diagonal elements equal to the reef-specific value 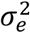 (and thus comparable in average magnitude of environmental variability to the other scenarios).

#### Estimating VPRSA from simulated community dynamics data

We tested the robustness of statistical inferences drawn from VPRSA by analyzing the simulated data described above. Following the approach used with the real data, we used the R package “poilog” to fit the bivariate Poisson-lognormal distributions to paired assemblage at different time lags, and then analysed the correlation coefficients for all of these pairs as functions of the time elapsed between them, to estimate the three parameters (*ρ*_0_, *ρ*_∞_, and *δ*) of the autocorrelation function (eq. 1) for all 40 reefs simultaneously using the nonlinear mixed-effects modelling approach employed in the main text analysis. Consistent with our main text analysis, we included random effects on 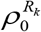 and 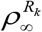 but not *δ* to avoid numerical instability of parameter estimates (Supplementary Text). From these estimates, we calculated the proportion of the total variance of log-abundance attributable to persistent intrinsic species differences, responses to environmental fluctuations, and demographic and sampling variance (overdispersion), as in the original analysis.

#### Testing the robustness of statistical inferences drawn from VPRSA

Once we had our reef-level variance component estimates, we then assessed the robustness of estimated proportional variance components to violations of model assumptions (scenarios ii through iv). For these simulations, we used average values of density dependence and environmental variance and covariance parameters (across species) for computing “expected” proportional variances (i.e., according to eq. 7 in the main text) in scenarios of varied intra- and inter-specific interactions (scenarios ii and iii) and environmental variances and covariances (scenarios iv and v). Thus, for example, for the case of variable intra-specific density-dependence, we compared the variance components obtained from analysis of the simulated data, with the theoretical expectation under the assumption that all species exhibited the average level of density dependence (i.e., the average δ to substitute δ in eq. 7 in the main text) for that simulation. Our goal is to determine whether the proportional variance estimates produced by applying the VPRSA approach, to simulated data that violate the model assumptions, produce estimates that are consistent with the overall relative importance of persistent species differences versus stochastic fluctuations in those simulated data.

In addition, we calculated, analytically, an alternative measure of theoretical expected proportional variances to take more explicit account of between-species heterogeneity, species interactions, and covariances in response to environmental fluctuations (termed the “Robust” predictions, below). We did this by exploiting general analytical solutions (*52*) for the environmental variance and variance in equilibrium abundance for the discrete time, stochastic, multivariate Gompertz model as follows:

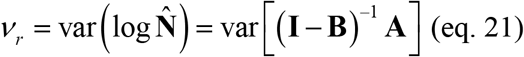

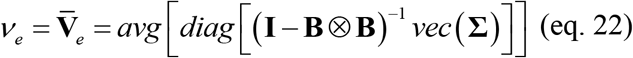

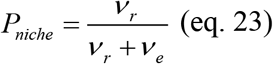

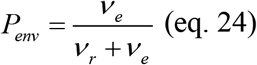

where *v_r_* is the among-species variance of equilibrium population sizes (on a logarithmic scale), and *v_e_* represents the average species-level variance of log-abundance due to environmental stochasticity. In (eq. 21), log **N̂** represents the species’ abundances at stationary or equilibrium states, “var” represents the variance operator, **B** is the interaction matrix (as per eq. 9), **A** is a vector of intrinsic growth rates (as per eq. 9). In (eq. 22), **V_e_** is the average of the diagonal of the environmental variance-covariance matrix at stationary states, “avg” represents the arithmetic mean function, and “diag” and “vec” are the diagonal and vectorization operators. The symbol Ä represents the Kronecker or tensor product. From equations 23-24, the two variance components are generalized to account for heterogeneity in intra-specific and inter-specific density dependences through the interaction matrix **B**, as well as heterogeneity in environmental variances and covariances through the environmental variance-covariance matrix **Σ**.

Importantly, under the assumptions of the *Baseline* scenario, the approximate measures of *v_r_* and *v_e_*above (eqs. 21-24) collapse to discrete-time analogous of Engen and colleagues’ functional forms of *V_r_* and *V_e_* (eq. 7 in the main text) (where the density-dependent parameter *δ* ≡ 1 − *b_ii_*). However, once species interactions or hterogeneity in density-dependence are incorporated (e.g., scenarios ii and iii), 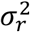 would cease to be directly proportional to the variance in species’ equilibrium log-abundances, so we would expect this modified version of *vr* (eq. 21) to better measure the relative importance of persistent niche structure than *Vr* from the original theory (eq. 7 in the main text). Similarly, in the presence of environmental covariances or heterogeneity in environmental variances among species (e.g., scenarios iv and v), the functional relationship between *Ve* in eq. 7 and the overall contribution of environmentally mediated population fluctuations to variance in species-abundances may also break down, rendering eq. 22 a more robust measure. Consequently, for scenarios (ii)-(v), we test estimated variance components from our fits against theoretical variance components calculated according to both the original theory (eq. 7 in the main text), and the generalized forms above (eqs. 21-24). However, as noted above, the model from which these generalized forms are derived (eqs. 21-22) omits demographic stochasticity; thus, if the contribution of *Vd* in eq. 7 is non-negligible, then analytically-calculated proportional variance components from eqs. 23-24 will be biased. Consequently, to maximize the comparability of these quantities *v_r_* and *v_e_* with the original VPRSA forms, we normalized the variance components of the former *V_r_* and *V_e_* (eq. 7) as follows:

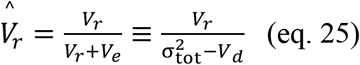

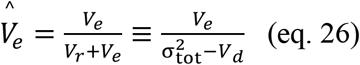

As expected, estimated variance components from data simulated according to our “baseline” scenario were highly consistent with the theoretical expectation. In Simulation Study Fig. 2, each point represents variance component estimates for one reef in one simulation. The colored line represents the theoretical expectation for the variance component due to persistent species differences (red), and environmental stochasticity (blue), and demographic and sampling variance (green). The black line is a smoothed fit to the simulated data. Thus, the discrepancy between the black line and the corresponding colored line represents the difference between the underlying trend in the simulated data, versus the theoretical expectation according to the original theoretical model. For this baseline scenario, there is a slight tendency to overestimate, by a few percentage points on average, the contribution of persistent species differences relative to stochastic fluctuations, particularly when the contribution of stochastic fluctuations is large. This discrepancy is not markedly increased by the incorporation of species interactions, interspecific variation in sensitivity to environmental fluctuations, or covariation between species in fluctuations, but is slightly larger in the presence of variation in intra-specific density-dependence (Simulation Study Fig. 3).

Our “robust approach” seems to better capture the behavior of the variance components than the original theory (Simulation Study Fig. 4 and Fig. 5). Specifically, the central tendency of the variance component estimates aligns much more closely with the variance components predicted by the more general theoretical model given by eqs 21-24 than it does to the predictions of the original theory (Simulation Study Figs. 4-5). To understand this, it is important to note that, conceptually, the variance attributable to “persistent species differences” is the variance in deterministic equilibrium population sizes (on a logarithmic scale). In Engen et al.’s (2002) original model, as in the baseline scenario, this is equal to 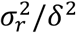in eq. 7 (i.e., *ρ*_∞_ in eq. 1) in the main text. However, when there is heterogeneity among species in intra-specific density dependence (i.e., the “varIntra” scenario; Simulation Study Fig. 4A-B) or species interactions (i.e., the “varInter” scenario; Simulation Study Fig. 4C-D), or when there are different species responses to environmental fluctuations (i.e., the “varEnv” and “varcovEnv” scenario; Simulation Study Fig. 5), this is no longer true, and thus the original analytical expectation (eq. 7 in the main text) itself may be less representative of the true variance components of relative species abundance. In this context, it is encouraging that the VPRSA estimates produced by analysis of the autocorrelation function align well with the “robust” formulation that takes the additional heterogeneity among species into account with the appropriate multivariate Gompertz expressions (eqs. 21-24) (Simulation Study Fig. 4 and Fig. 5). In other words, the estimated proportional variance component for persistent species differences appears to provide a very good estimate of the relative amount of the variance in species’ log-abundances that is due to variance in their long-term equilibrium values, relative to the variance due to their temporal fluctuations. Notably, the modified expressions (eqs. 21-24) also perform better in the baseline scenario (i.e., when species differ only in intrinsic growth rates), compared to the original analytical expressions (eq. 7 in the main text) (Simulation Study Fig. 6), which suggests that some of the discrepancy may be due to the discrete-time nature of the simulations (since the original theoretical model was developed for continuous-time dynamics, whereas the theoretical model underpinning our “robust approach” is formulated in discrete time).

These supplementary simulation results suggest that proportional variance estimates from VPRSA provide robust information about the relative importance of persistent species differences (determinism) versus stochastic environmental fluctuations in shaping patterns of commonness and rarity among species, even when key simplifying assumptions about community dynamics made by the original theory are violated. We conclude from this that VPRSA may well be much more broadly applicable than previously realized, particularly when the variance component due to persistent species differences is conceptualized as representing the proportional variance in species abundances due to differences in their long-term mean abundances (as in our derivations from the more general theory), rather than specifically to differences in their intrinsic growth rates (as in the original theory used to derive the variance components). Of course, our simulation study cannot be a comprehensive exploration of the robustness of this approach to all possible assumption violations. However, the robustness that we have identified suggests that VPRSA is more robust than one might have assumed, given the original model of community dynamics that inspired it.

### Simulation Study Figs. 1 to 6

**Simulation Study Fig. 1.**
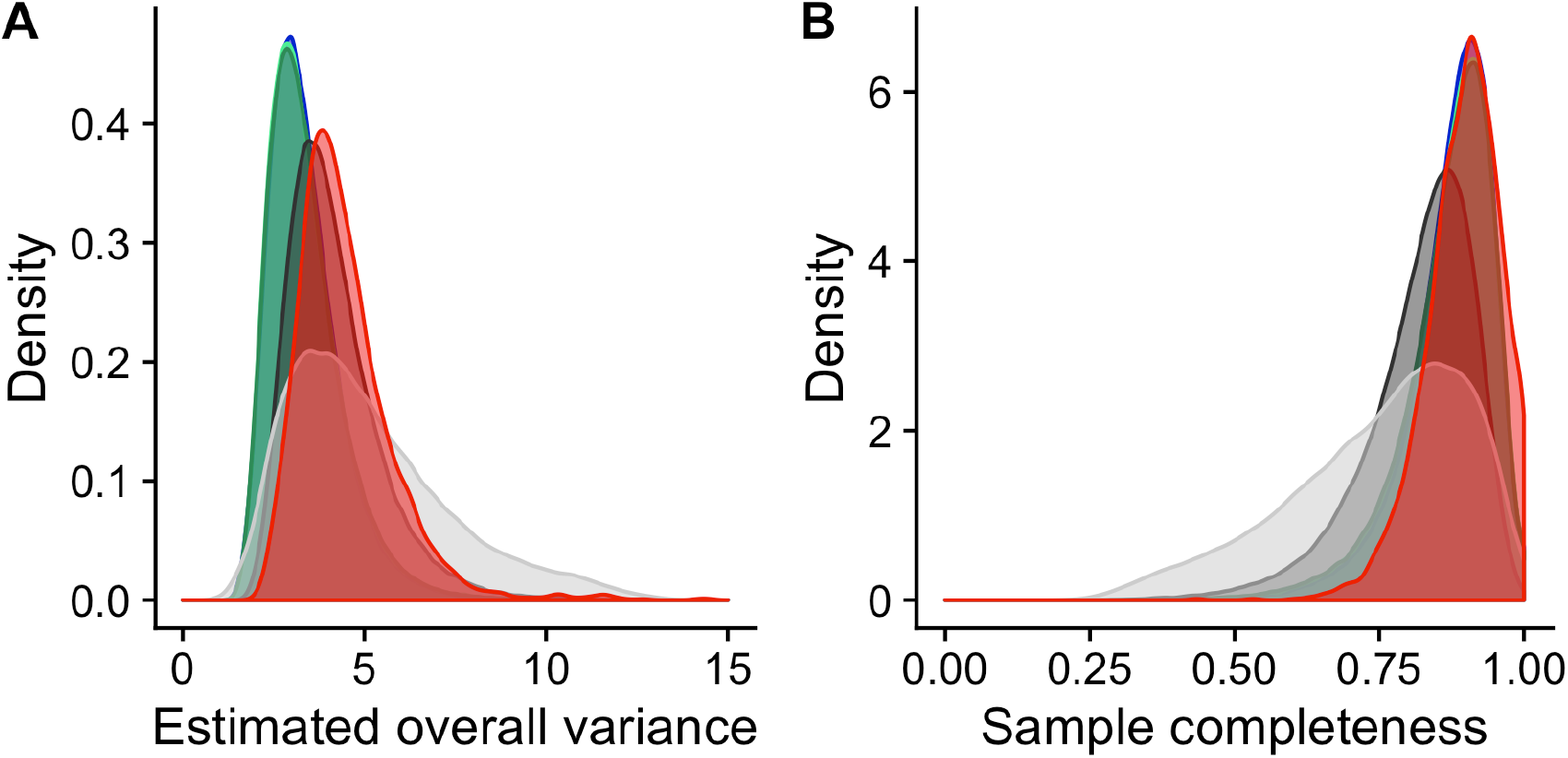
(**A**) Density distribution of estimated overall variance of Poisson-lognormal species-abundance distributions. (**B**) Density distribution of sample completeness measured as the ratio of observed (sampled) species richness to estimated species richness from Poisson-lognormal fits. Curves are probability density distributions from empirical and simulated data. Red curves represent the empirical LTMP data. Green curves (light green, intermediate, and dark green) are ‘*baseline*’,’*varcovEnv*’, and ‘*varEnv*’ scenarios, respectively. Light- and dark-gray curves are ‘*varIntra*’ and ‘*varInter*’ scenarios, respectively.

**Simulation Study Fig. 2.**
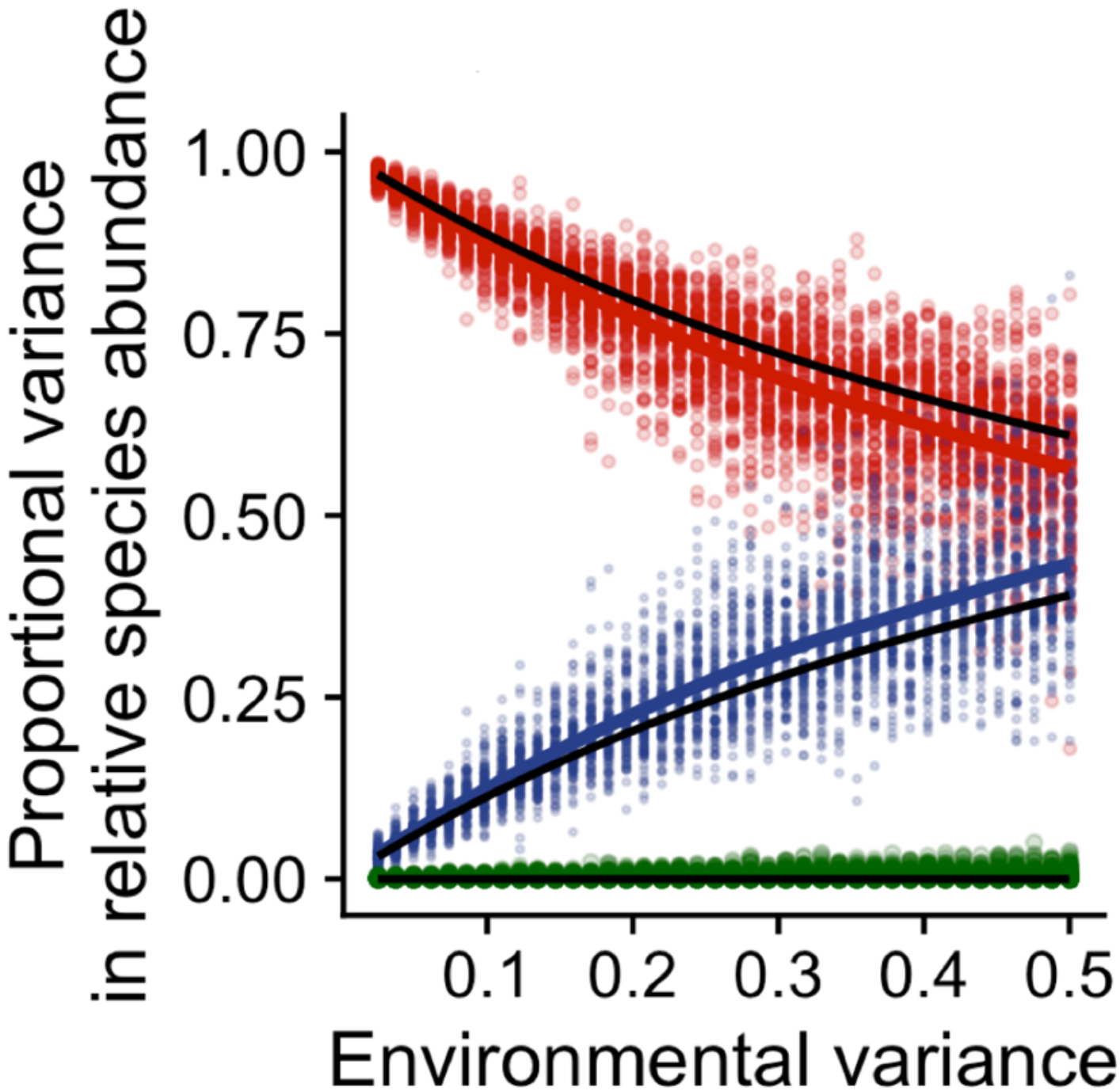
Relationships between true environmental variance and VPRSA-estimated variance components of persistent species differences, environmental stochasticity, and a combination of demographic or sampling stochasticity for the “Baseline” scenario. Red, blue, and green colors represent the VPRSA-estimated proportional variance driven by persistent species differences (niche structure), environmental stochasticity, and demographic or sampling stochasticity, respectively. Each point, irrespective of color, represents one simulated time series for one spatial replicate (reef) (i.e., for each color, n = 40 spatial replicates with varied environmental variance × 100 simulations = 400 points). All community dynamics data (points) are simulated from the baseline scenario. Red, blue, and green colored lines represent the kernel smoothing of proportional variance estimates, respectively, obtained using local polynomial regression fitting. Black lines represent the analytical predictions of Engen et al. 2002 (eq. 7 in the main text) using the true parameters from the simulations.

**Simulation Study Fig. 3.**
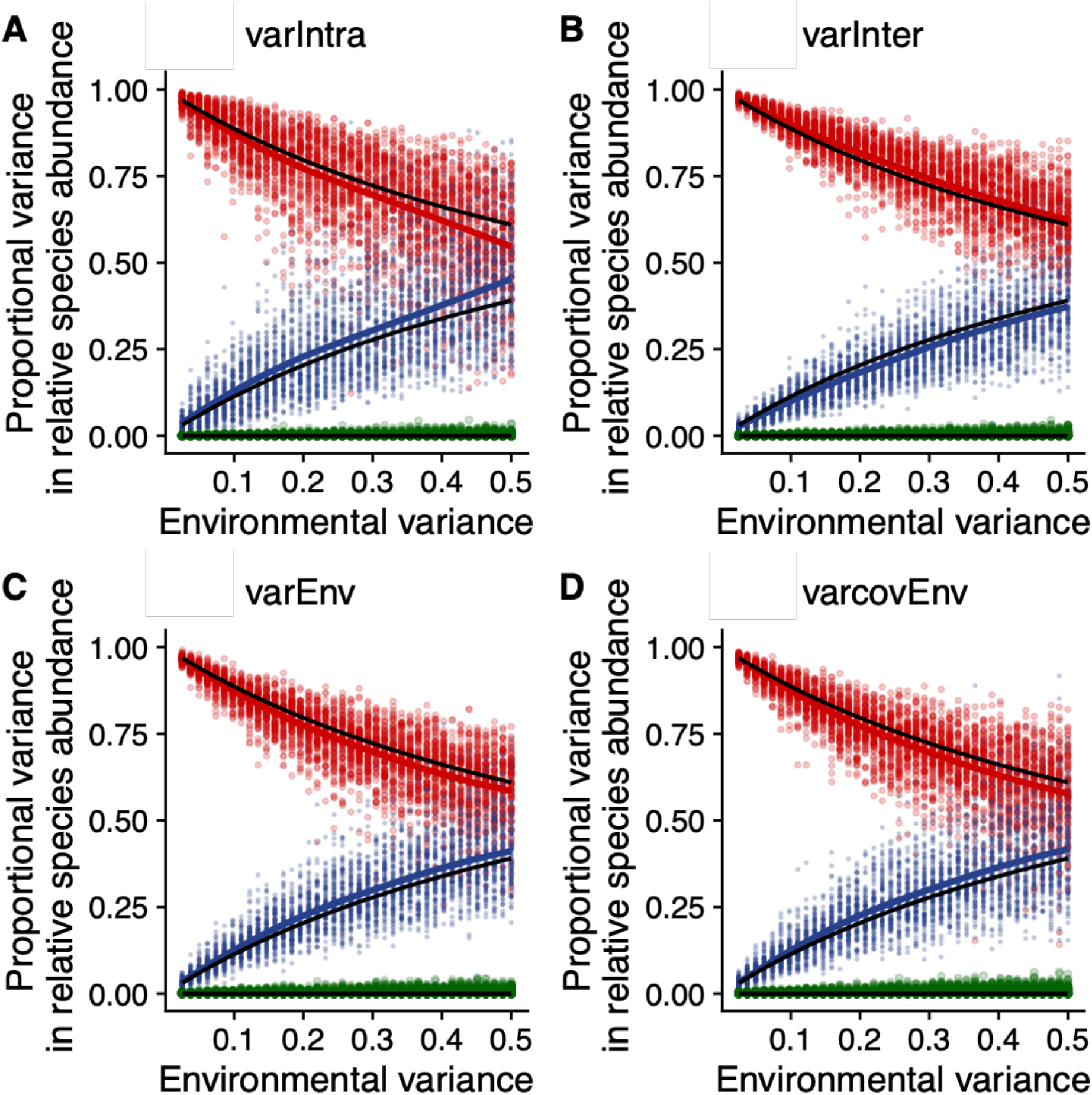
Relationships between true environmental variance and estimates of variance components under different community dynamics scenarios. Red, blue, and green colors represent the proportional variance components due to persistent species differences, environmental stochasticity, and demographic or sampling stochasticity. Colored (red, blue, and green) lines represent the kernel smoothing of proportional variance estimates, respectively, obtained using local polynomial regression fitting. Black lines represent the analytical prediction of Engen et al. 2002 (eq. 7 in the main text) using the true parameters from the simulations. (**A**) The “*varIntra*” scenario, which includes species differences in intra-specific density dependence. (**B**) The “*varInter*” scenario, which includes species differences in inter-specific density dependence. (**C**) The “*varEnv*” scenario, which includes species differences in the magnitude of environmental variance. (**D**) The “*varcovEnv*” scenario, in which species’ responses to environmental fluctuations covary.

**Simulation Study Fig. 4.**
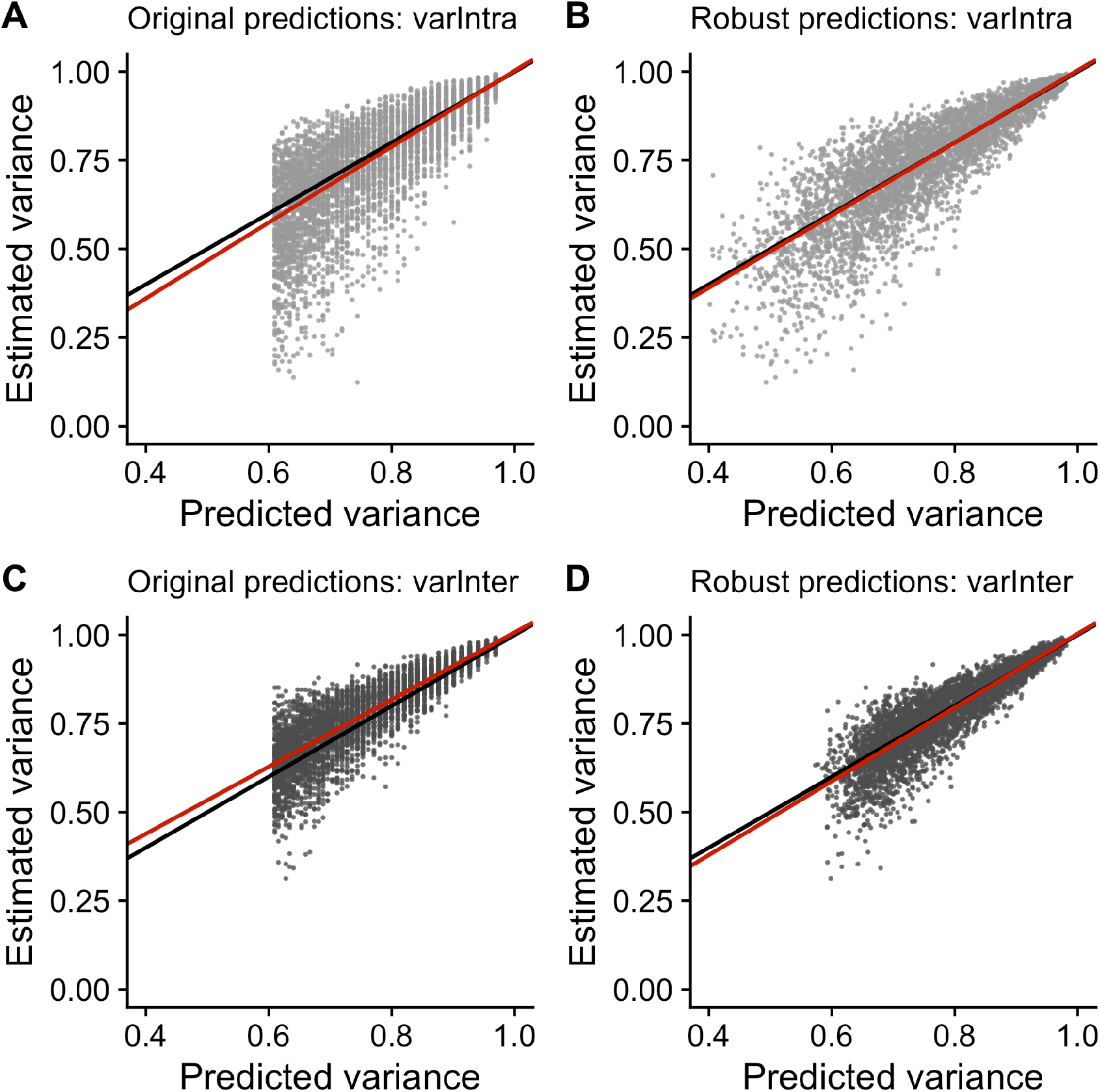
Relationships between variance estimate of persistent species differences and its analytical prediction. The black line is the unity line indicating perfect agreement between VPRSA estimates and analytical predictions. The red line is a quantile regression through the median of the VPRSA estimates of variance components of persistent species differences. (**A**, **C**) The relationship between VPRSA estimates and the original analytical prediction of Engen et al. 2002 (eq. 7 in the main text), and (**B**, **D**) the relationship between VPRSA estimates and the generalized analytical prediction from the discrete-time multivariate Gompertz model (eqs. 21-24) under the (**A**, **B**) “varIntra” and (**C**, **D**) “varInter” community dynamics scenarios.

**Simulation Study Fig. 5.**
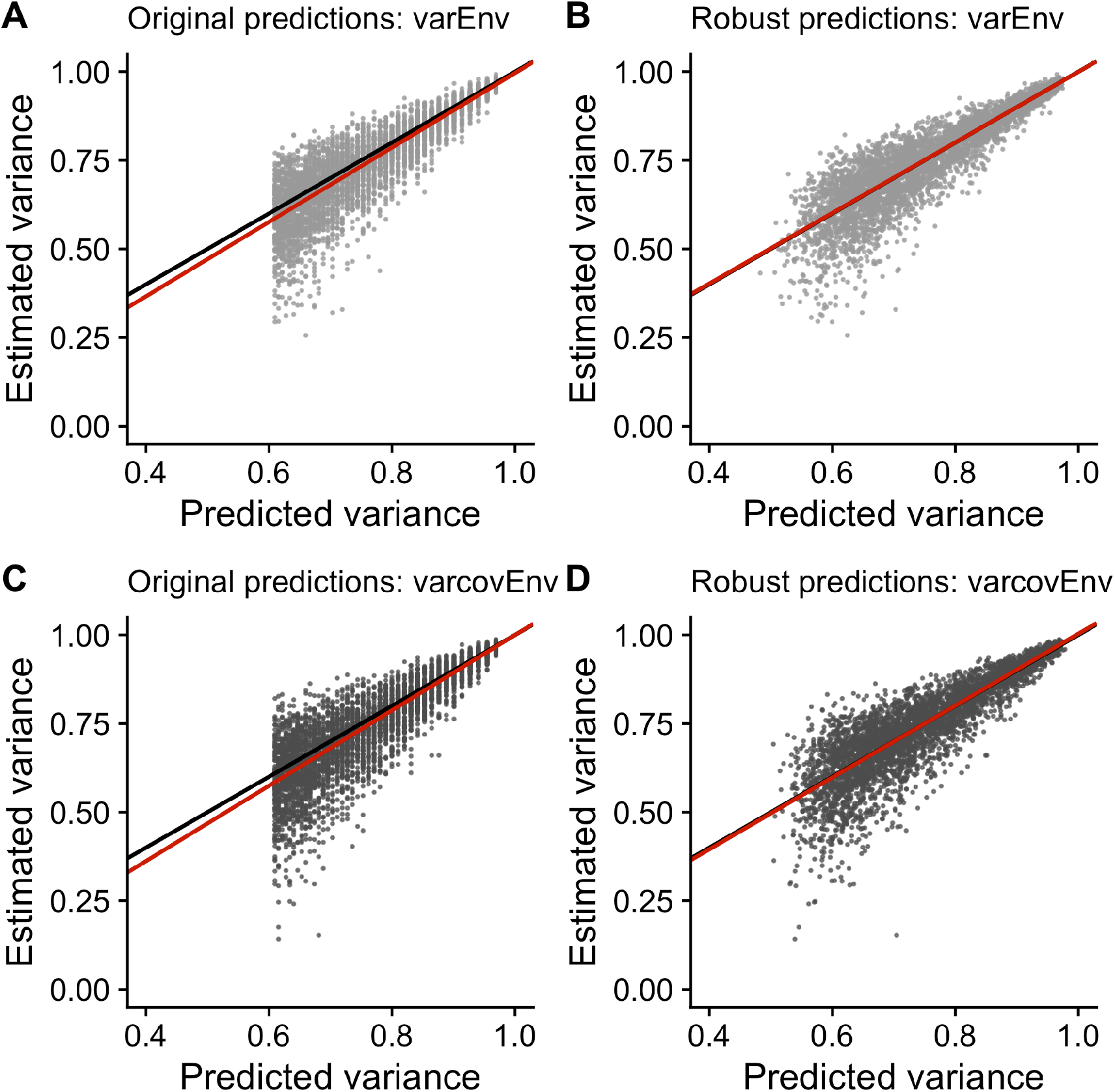
Relationships between variance estimate of persistent species differences and its analytical prediction. The black line is the unity line indicating perfect agreement between VPRSA estimates and analytical predictions. The red line is a quantile regression through the median of the VPRSA estimates of variance components of persistent species differences. (**A**, **C**) The relationship between VPRSA estimates and the original analytical prediction of Engen et al. 2002 (eq. 7 in the main text), and (**B**, **D**) the relationship between VPRSA estimates and the generalized analytical prediction from the discrete-time multivariate Gompertz model (eqs. 21-24) under the (**A**, **B**) “varEnv” and (**C**, **D**) “varcovEnv” community dynamics scenarios.

**Simulation Study Fig. 6.**
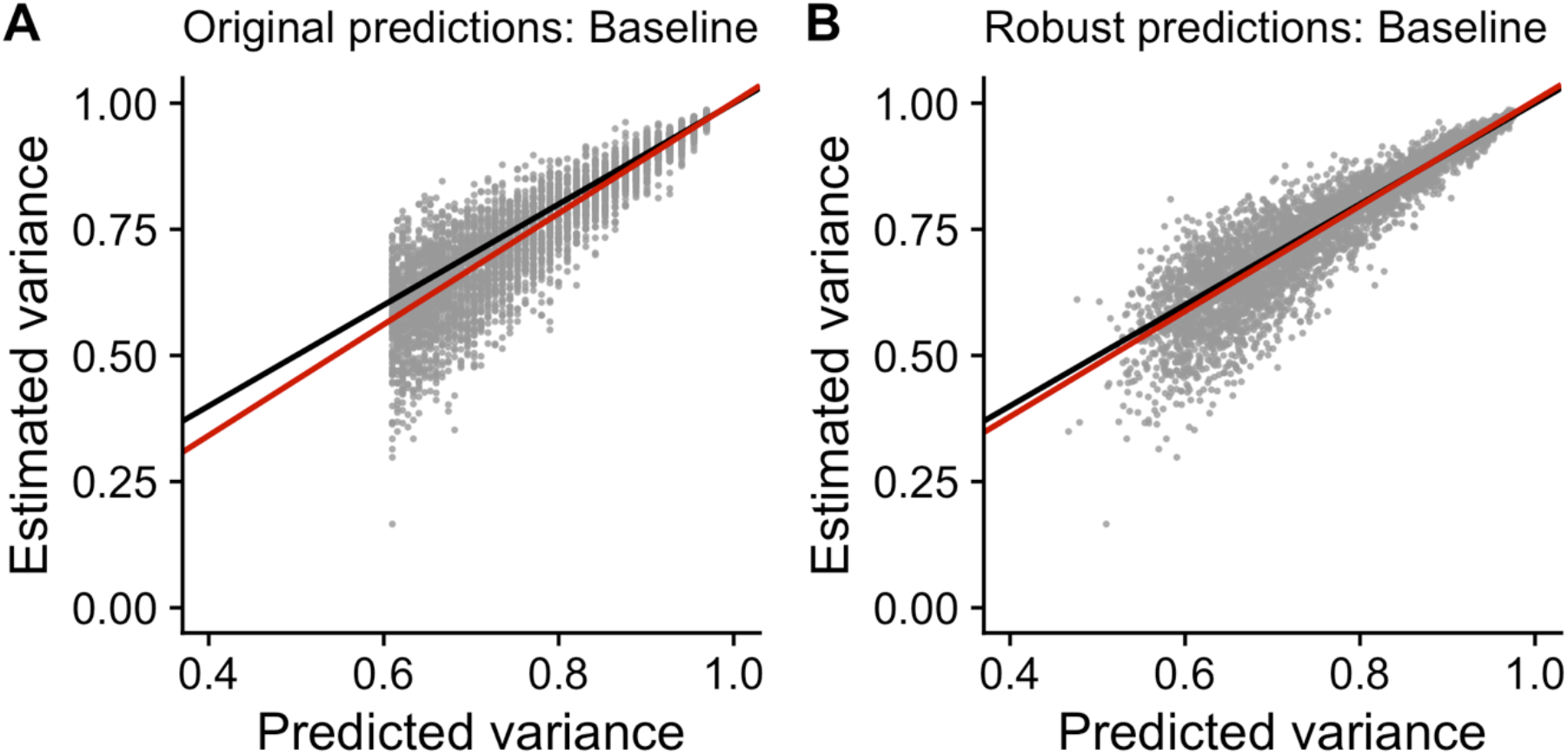
Relationships between variance estimate of persistent species differences and its analytical prediction under baseline scenario. The black line is the unity line indicating perfect agreement between VPRSA estimates and analytical predictions. The red line is a quantile regression through the median of the VPRSA estimates of variance components of persistent species differences. (**A**) The relationship between VPRSA estimates and the original analytical prediction of Engen et al. 2002 (eq. 7 in the main text), and (**B**) the relationship between VPRSA estimates and the generalized analytical prediction from the discrete-time multivariate Gompertz model (eqs. 21-24).

## Notes

### Competing Interest Statement

The authors have declared no competing interest.

## References and Notes

1. J. H. Connell, Diversity in tropical rain forests and coral reefs. Science. 199, 1302–1310 (1978).

2. P. L. Chesson, R. R. Warner, Environmental variability promotes coexistence in lottery competitive systems. Am. Nat. 117, 923–943 (1981).

3. C. Syms, G. P. Jones, Disturbance, habitat structure, and the dynamics of a coral-reef fish communtiy. Ecology. 81, 2714–2729 (2000).

4. G. De’ath, K. E. Fabricius, H. Sweatman, M. Puotinen, The 27-year decline of coral cover on the Great Barrier Reef and its causes. Proc. Natl. Acad. Sci. U. S. A. 109, 17995–17999 (2012).

5. T. P. Hughes, J. T. Kerry, A. H. Baird, S. R. Connolly, A. Dietzel, C. M. Eakin, S. F. Heron, A. S. Hoey, M. O. Hoogenboom, G. Liu, M. J. McWilliam, R. J. Pears, M. S. Pratchett, W. J. Skirving, J. S. Stella, G. Torda, Global warming transforms coral reef assemblages. Nature. 556, 492–496 (2018).

6. T. P. Hughes, M. L. Barnes, D. R. Bellwood, J. E. Cinner, G. S. Cumming, J. B. C. Jackson, J. Kleypas, I. A. van de Leemput, J. M. Lough, T. H. Morrison, S. R. Palumbi, E. H. van Nes, M. Scheffer, Coral reefs in the Anthropocene. Nature. 546, 82–90 (2017).

7. E. S. Darling, T. R. McClanahan, I. M. Côté, Life histories predict coral community disassembly under multiple stressors. Glob. Chang. Biol. 19, 1930–1940 (2013).

8. S. K. Wilson, N. A. J. Graham, M. S. Pratchett, G. P. Jones, N. V. C. Polunin, Multiple disturbances and the global degradation of coral reefs: Are reef fishes at risk or resilient? Glob. Chang. Biol. 12 (2006), pp. 2220–2234.

9. M. Gouezo, Y. Golbuu, K. Fabricius, D. Olsudong, G. Mereb, V. Nestor, E. Wolanski, P. Harrison, C. Doropoulos, Drivers of recovery and reassembly of coral reef communities. Proc. R. Soc. B. 286, 20182908 (2019).

10. T. Lamy, R. Galzin, M. Kulbicki, T. L. De Loma, J. Claudet, Three decades of recurrent declines and recoveries in corals belie ongoing change in fish assemblages. Coral Reefs. 35, 293–302 (2016).

11. J. M. Pandolfi, J. B. C. Jackson, Ecological persistence interrupted in Caribbean coral reefs. Ecol. Lett. 9, 818–826 (2006).

12. P. J. Edmunds, M. Adjeroud, M. L. Baskett, I. B. Baums, A. F. Budd, R. C. Carpenter, N. S. Fabina, T.-Y. Fan, E. C. Franklin, K. Gross, Persistence and change in community composition of reef corals through present, past, and future climates. PLoS One. 9, e107525 (2014).

13. W. A. DiMichele, A. K. Behrensmeyer, T. D. Olszewski, C. C. Labandeira, J. M. Pandolfi, S. L. Wing, R. Bobe, Long-term stasis in ecological assemblages: evidence from the fossil record. Annu. Rev. Ecol. Evol. Syst. 35, 285–322 (2004).

14. S. P. Hubbell, The unified neutral theory of biodiversity and biogeography (MPB-32) (Princeton University Press, 2001).

15. M. Dornelas, S. R. Connolly, T. P. Hughes, Coral reef diversity refutes the neutral theory of biodiversity. Nature. 440, 80–82 (2006).

16. I. Volkov, J. R. Banavar, S. P. Hubbell, A. Maritan, Patterns of relative species abundance in rainforests and coral reefs. Nature. 450, 45 (2007).

17. M. Bode, S. R. Connolly, J. M. Pandolfi, Species differences drive nonneutral structure in pleistocene coral communities. Am. Nat. 180, 577–88 (2012).

18. S. R. Connolly, M. A. MacNeil, M. J. Caley, N. Knowlton, E. Cripps, M. Hisano, L. M. Thibaut, B. D. Bhattacharya, L. Benedetti-Cecchi, R. E. Brainard, A. Brandt, F. Bulleri, K. E. Ellingsen, S. Kaiser, I. Kröncke, K. Linse, E. Maggi, T. D. O’Hara, L. Plaisance, G. C. B. Poore, S. K. Sarkar, K. K. Satpathy, U. Schückel, A. Williams, R. S. Wilson, Commonness and rarity in the marine biosphere. Proc. Natl. Acad. Sci. U. S. A. 111, 8524–8529 (2014).

19. . R. Lande, S. Engen, B. Saether, Stochastic population dynamics in ecology and conservation (Oxford University Press on Demand, 2003).

20. A. S. Hoey, D. R. Bellwood, Cross-shelf variation in browsing intensity on the Great Barrier Reef. Coral Reefs. 29, 499–508 (2010).

21. A. Cheal, M. Emslie, I. Miller, H. Sweatman, The distribution of herbivorous fishes on the Great Barrier Reef. Mar. Biol. 159, 1143–1154 (2012).

22. V. Y. Y. Lam, M. Chaloupka, A. Thompson, C. Doropoulos, P. J. Mumby, Acute drivers influence recent inshore Great Barrier Reef dynamics. Proc. R. Soc. B. 285, 20182063 (2018).

23. C. Mellin, C. J. A. Bradshaw, M. G. Meekan, M. J. Caley, Environmental and spatial predictors of species richness and abundance in coral reef fishes. Glob. Ecol. Biogeogr. 19, 212–222 (2010).

24. A. J. Cheal, M. A. MacNeil, M. J. Emslie, H. Sweatman, The threat to coral reefs from more intense cyclones under climate change. Glob. Chang. Biol. 23, 1511–1524 (2017).

25. M. S. Pratchett, C. F. Caballes, J. C. Wilmes, S. Matthews, C. Mellin, H. Sweatman, L. E. Nadler, J. Brodie, C. A. Thompson, J. Hoey, Thirty years of research on crown-of-thorns starfish (1986–2016): scientific advances and emerging opportunities. Diversity. 9, 41 (2017).

26. T. P. Hughes, J. T. Kerry, M. Álvarez-Noriega, J. G. Álvarez-Romero, K. D. Anderson, A. H. Baird, R. C. Babcock, M. Beger, D. R. Bellwood, R. Berkelmans, T. C. Bridge, I. R. Butler, M. Byrne, N. E. Cantin, S. Comeau, S. R. Connolly, G. S. Cumming, S. J. Dalton, G. Diaz-Pulido, C. M. Eakin, W. F. Figueira, J. P. Gilmour, H. B. Harrison, S. F. Heron, A. S. Hoey, J.-P. A. Hobbs, M. O. Hoogenboom, E. V. Kennedy, C. Kuo, J. M. Lough, R. J. Lowe, G. Liu, M. T. McCulloch, H. A. Malcolm, M. J. McWilliam, J. M. Pandolfi, R. J. Pears, M. S. Pratchett, V. Schoepf, T. Simpson, W. J. Skirving, B. Sommer, G. Torda, D. R. Wachenfeld, B. L. Willis, S. K. Wilson, Global warming and recurrent mass bleaching of corals. Nature. 543, 373–377 (2017).

27. T. P. Hughes, K. D. Anderson, S. R. Connolly, S. F. Heron, J. T. Kerry, J. M. Lough, A. H. Baird, J. K. Baum, M. L. Berumen, T. C. Bridge, D. C. Claar, C. M. Eakin, J. P. Gilmour, N. A. J. Graham, H. Harrison, J.-P. A. Hobbs, A. S. Hoey, M. Hoogenboom, R. J. Lowe, M. T. McCulloch, J. M. Pandolfi, M. Pratchett, V. Schoepf, G. Torda, S. K. Wilson, Spatial and temporal patterns of mass bleaching of corals in the Anthropocene. Science. 359, 80–83 (2018).

28. M. Dornelas, N. J. Gotelli, B. McGill, H. Shimadzu, F. Moyes, C. Sievers, A. E. Magurran, Assemblage time series reveal biodiversity change but not systematic loss. Science. 344, 296–299 (2014).

29. S. A. Blowes, S. R. Supp, L. H. Antão, A. Bates, H. Bruelheide, J. M. Chase, F. Moyes, A. Magurran, B. McGill, I. H. Myers-Smith, M. Winter, A. D. Bjorkman, D. E. Bowler, J. E. K. Byrnes, A. Gonzalez, J. Hines, F. Isbell, H. P. Jones, L. M. Navarro, P. L. Thompson, M. Vellend, C. Waldock, M. Dornelas, The geography of biodiversity change in marine and terrestrial assemblages. Science. 366, 339–345 (2019).

## Supplementary References

30. . H. Sweatman, S. Burgess, A. Cheal, G. Coleman, S. Delean, M. Emslie, A. McDonald, I. Miller, K. Osborne, A. Thompson, Long-term monitoring of the Great Barrier Reef. Status Report Number 7 (2005).

31. S. Engen, R. Lande, T. Walla, P. J. DeVries, Analyzing spatial structure of communities using the two-dimensional Poisson Lognormal species abundance model. Am. Nat. 160, 60–73 (2002).

32. . R. Lande, S. Engen, B. Saether, Stochastic population dynamics in ecology and conservation (Oxford University Press on Demand, 2003).

33. B. Sæther, S. Engen, V. Grøtan, Species diversity and community similarity in fluctuating environments: parametric approaches using species abundance distributions. J. Anim. Ecol. 82, 721–738 (2013).

34. S. R. Connolly, M. A. MacNeil, M. J. Caley, N. Knowlton, E. Cripps, M. Hisano, L. M. Thibaut, B. D. Bhattacharya, L. Benedetti-Cecchi, R. E. Brainard, A. Brandt, F. Bulleri, K. E. Ellingsen, S. Kaiser, I. Kröncke, K. Linse, E. Maggi, T. D. O’Hara, L. Plaisance, G. C. B. Poore, S. K. Sarkar, K. K. Satpathy, U. Schückel, A. Williams, R. S. Wilson, Commonness and rarity in the marine biosphere. Proc. Natl. Acad. Sci. U. S. A. 111, 8524–8529 (2014).

35. L. M. Thibaut, S. R. Connolly, H. Sweatman, Diversity and stability of herbivorous fishes on coral reefs. Ecology. 93, 891–901 (2012).

36. E. B. Solbu, O. H. Diserud, J. A. Kålås, S. Engen, Heterogeneity among species and community dynamics—Norwegian bird communities as a case study. Ecol. Modell. 388, 13–23 (2018).

37. S. Engen, K. Aagaard, T. Bongard, Disentangling the effects of heterogeneity, stochastic dynamics and sampling in a community of aquatic insects. Ecol. Modell. 222, 1387–1393 (2011).

38. B. Smith, J. B. Wilson, A consumer’s guide to evenness indices. Oikos, 70–82 (1996).

39. S. Greven, T. Kneib, On the behaviour of marginal and conditional AIC in linear mixed models. Biometrika. 97, 773–789 (2010).

40. C. Wager, F. Vaida, G. Kauermann, Model selection for penalized spline smoothing using Akaike information criteria. Aust. N. Z. J. Stat. 49, 173–190 (2007).

41. S. A. Matthews, C. Mellin, A. MacNeil, S. F. Heron, W. Skirving, M. Puotinen, M. J. Devlin, M. Pratchett, High-resolution characterization of the abiotic environment and disturbance regimes on the Great Barrier Reef, 1985–2017. Ecology. 100, e02574 (2019).

42. B. J. McGill, R. S. Etienne, J. S. Gray, D. Alonso, M. J. Anderson, H. K. Benecha, M. Dornelas, B. J. Enquist, J. L. Green, F. He, Species abundance distributions: moving beyond single prediction theories to integration within an ecological framework. Ecol. Lett. 10, 995–1015 (2007).

43. R. Mac Nally, J. R. Thomson, W. J. Kimmerer, F. Feyrer, K. B. Newman, A. Sih, W. A. Bennett, L. Brown, E. Fleishman, S. D. Culberson, Analysis of pelagic species decline in the upper San Francisco Estuary using multivariate autoregressive modeling (MAR). Ecol. Appl. 20, 1417–1430 (2010).

44. J. Knape, P. de Valpine, Are patterns of density dependence in the Global Population Dynamics Database driven by uncertainty about population abundance? Ecol. Lett. 15, 17–23 (2012).

45. L. M. Thibaut, S. R. Connolly, Hierarchical modeling strengthens evidence for density dependence in observational time series of population dynamics. Ecology. 101, e02893 (2020).

46. R. A. Myers, K. G. Bowen, N. J. Barrowman, Maximum reproductive rate of fish at low population sizes. Can. J. Fish. Aquat. Sci. 56, 2404–2419 (1999).

47. M. C. Bonin, L. Boström-Einarsson, P. L. Munday, G. P. Jones, The prevalence and importance of competition among coral reef fishes. Annu. Rev. Ecol. Evol. Syst. 46, 169–190 (2015).

48. R. P. Freckleton, O. T. Lewis, Pathogens, density dependence and the coexistence of tropical trees. Proc. R. Soc. B Biol. Sci. 273, 2909–2916 (2006).

49. G. Gellner, K. S. McCann, Consistent role of weak and strong interactions in high-and low-diversity trophic food webs. Nat. Commun. 7, 1–7 (2016).

50. M. Loreau, From populations to ecosystems: Theoretical foundations for a new ecological synthesis (MPB-46) (Princeton University Press, 2010).

51. S. Engen, R. Lande, Population dynamic models generating the lognormal species abundance distribution. Math. Biosci. 132, 169–183 (1996).

52. A. R. Ives, B. Dennis, K. L. Cottingham, S. R. Carpenter, Estimating community stability and ecological interactions from time-series data. Ecol. Monogr. 73, 301–330 (2003).

53. C. Boettiger, From noise to knowledge: how randomness generates novel phenomena and reveals information. Ecol. Lett. 21, 1255–1267 (2018).

